# Enhancer-promoter hubs organize transcriptional networks promoting oncogenesis and drug resistance

**DOI:** 10.1101/2024.07.02.601745

**Authors:** Brent S. Perlman, Noah Burget, Yeqiao Zhou, Gregory W. Schwartz, Jelena Petrovic, Zora Modrusan, Robert B. Faryabi

## Abstract

Recent advances in high-resolution mapping of spatial interactions among regulatory elements support the existence of complex topological assemblies of enhancers and promoters known as enhancer-promoter hubs or cliques. Yet, organization principles of these multi-interacting enhancer-promoter hubs and their potential role in regulating gene expression in cancer remains unclear. Here, we systematically identified enhancer-promoter hubs in breast cancer, lymphoma, and leukemia. We found that highly interacting enhancer-promoter hubs form at key oncogenes and lineage-associated transcription factors potentially promoting oncogenesis of these diverse cancer types. Genomic and optical mapping of interactions among enhancer and promoter elements further showed that topological alterations in hubs coincide with transcriptional changes underlying acquired resistance to targeted therapy in T cell leukemia and B cell lymphoma. Together, our findings suggest that enhancer-promoter hubs are dynamic and heterogeneous topological assemblies with the potential to control gene expression circuits promoting oncogenesis and drug resistance.

## INTRODUCTION

Genome spatial organization facilitates enhancer-promoter communication, which is crucial for control of oncogenic transcriptional programs ^1, 2^. Emerging evidence from studies of cancer genome topology supports that multiple enhancers and promoters can spatially coalesce, forming topological assemblies that are variably referred to as enhancer-promoter hubs or cliques ^3, 4^. Nevertheless, the fundamental properties of these topological assemblies and their potential role in promoting oncogenesis remain unclear.

Investigation of oncogenic enhancer-promoter hubs has unique potential to advance our understanding of cancer given that enhancer dysregulation is a key hallmark of oncogenesis ^5^. Furthermore, current models have yet to fully grasp how distal enhancers exert their regulatory functions across large genomic distances. Genome topology, which is organized at various length scales from megabase-scale compartments and topologically associating domains (TADs) to fine-scale chromatin loops, contributes to spatial positioning of enhancers and their target promoters, influencing their activity and specificity ^6, 7, 8, 9^. Given that the number of active enhancers is 2-3 times more than active genes ^10^, it is often possible that multiple enhancers control the expression of a single gene, giving rise to complex enhancer regulatory circuits ^11, 12, 13^. Although chromatin interaction data alone cannot capture the complexity of potential multi-enhancer regulation, its integration with chromatin activity datasets at a few oncogenes revealed that multiple distal enhancers can spatially cluster with promoters to form enhancer-promoter hubs in cancer genomes ^3, 14, 15, 16, 17^. More recent studies have demonstrated that enhancer-promoter hubs facilitate enhancer cooperativity and target specificity to control gene expression dosage ^12, 14, 18^. Despite these advances, a systematic understanding of enhancer-promoter hub prevalence, organization principles, and regulatory importance in mediating oncogenic enhancer function is lacking.

Here, we systematically identified enhancer-promoter hubs in T cell acute lymphoblastic leukemia (T-ALL), mantle cell lymphoma (MCL), and triple negative breast cancer (TNBC) to elucidate prevalence and organization principles of these topological assemblies across diverse cancer types. Examination of enhancer-promoter hubs revealed that they are ubiquitous and distinct from TADs and super-enhancers. Study of T-ALL, MCL and TNBC enhancer-promoter interactions further showed that hubs are heterogeneous with asymmetric distribution of interactions among enhancers and promoters. Notably, a small subset of enhancer-promoter hubs was hyperinteracting, exhibiting exceptionally high spatial interactivity among constituent enhancer and promoter elements. We demonstrated that hyperinteracting hubs were uniquely enriched for transcription, predominantly formed around transcription factors and coregulators, and were more lineage associated than regular (i.e. non-hyperinteracting) hubs.

To further substantiate the structure-function relationship of enhancer-promoter hubs, we examined their reorganization in Notch inhibitor resistant T-ALL and Bruton’s tyrosine kinase (BTK) inhibitor resistant MCL. Population-based and single-cell resolution chromatin mapping studies revealed the role of enhancer-promoter hub reorganization in setting gene expression programs permissive to Notch inhibitor and BTK inhibitor resistance in T-ALL and MCL, respectively. Together, our data suggest that enhancer-promoter hub formation is an epigenetic mechanism which is potentially hijacked by cancer cells to set gene expression programs promoting oncogenesis and drug resistance.

## RESULTS

### Interactions among enhancers and promoters are asymmetrically distributed in T leukemic cells

Complex interactions among enhancers and promoters measured by high-resolution chromatin conformation capture assays such as *in-situ* Hi-C or HiChIP can be conceptualized as a network of connected nodes within nuclear space and modeled using undirected graph mathematical abstraction^14, 19^. To detect groups of highly interacting enhancers and promoters, known as enhancer-promoter hubs or cliques, from the graph of frequently interacting enhancers and promoters, we leveraged an efficient implementation of divisive hierarchical spectral clustering (see Methods) ^20^. Using global information about the enhancer-enhancer, enhancer-promoter, and promoter-promoter interactions embedded in the interactivity graph, our clustering approach identifies a hierarchy of densely interacting enhancer and promoter groups with high intra-group and sparse inter-group interactions (**Fig. 1a**). Notably, our implementation of divisive hierarchical spectral clustering has tunable parameters (see Methods), enabling identification of hubs with granularity that matches user preferences. Given that enhancer-promoter hubs are dually defined by regulatory element composition and spatial organization, we hypothesized that these topological assemblies organize transcription across cancer genomes.

**Figure 1:**
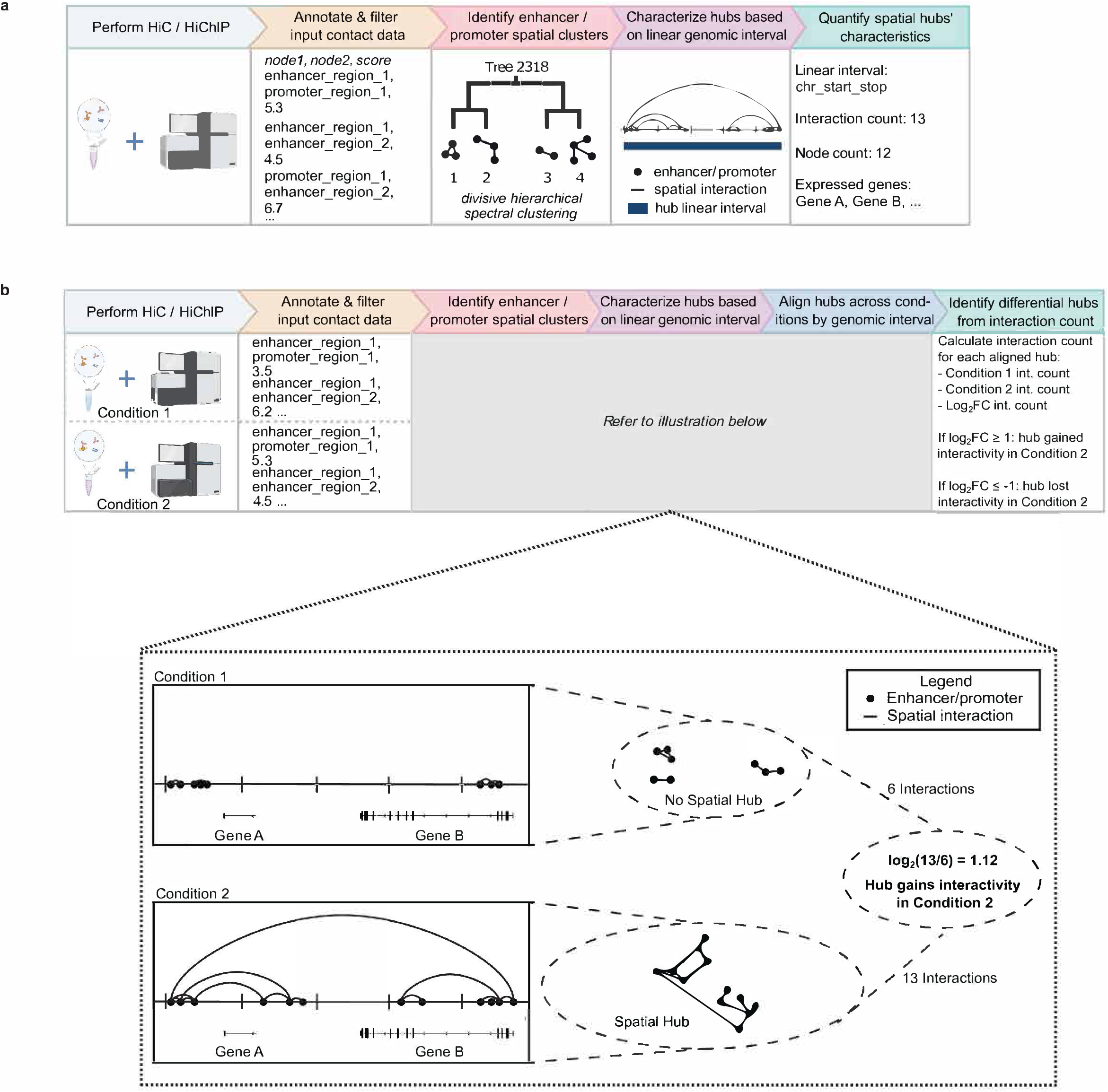
Enhancer-promoter hubs are systematically identified by divisive hierarchical spectral clustering of an undirected interactivity graph of enhancers and promoters. a: Schematic depicting the process of identifying enhancer-promoter hubs from raw chromatin conformation capture data. First, the interactivity graph of enhancer and promoter elements is created, where nodes are enhancers and promoters defined from H3K27ac ChIP-seq and RNA-seq measurements, respectively. These regulatory element nodes are connected by pairwise spatial interactions defined from normalized and filtered Hi-C or SMC1 HiChIP valid interactions. Next, an efficient matrix-free divisive hierarchical spectral clustering algorithm is used to partition the enhancer-promoter interactivity graph into spatial clusters. Clusters are then characterized by their contiguous linear genomic intervals from their most upstream spatially interacting regulatory element to their most downstream spatially interacting regulatory element to form enhancer-promoter hubs. These hubs can be ranked based on their interaction count, enhancer/promoter number, and the expressed genes contained within their linear genomic intervals. Input contact data depicted is for illustrative purposes only. b: Schematic depicting the identification of differential enhancer-promoter hubs between two conditions based on within-hub total interactivity between enhancers and/or promoters. Top: enhancer-promoter hubs separately identified in each condition are combined into a union set of hubs based on their linear genomic coordinates. Note that the input contact data depicted is for illustrative purposes only. Spatial enhancer hubs with markedly differential interactivity are identified based on log_2_ fold change of interaction count between the two conditions. Bottom: diagram of an illustrative differential hub. Bottom left depicts this hub on the linear genome where Hi-C valid contacts (arcs) connect enhancers and promoters (circle nodes) in condition 1 (upper) and condition 2 (lower) cells. Bottom right illustrates a simplified, potential 3D rendering of this hub in each condition, demonstrating that cells in condition 2 gain more than two-fold interactivity at this locus to form a (differential) spatial hub.

To evaluate this hypothesis, we first identified and characterized enhancer-promoter hubs using multi-omic data from DND41 T-ALL ^21^. From Hi-C data, 1,377 hubs with an average size of roughly 414 Kb were identified across the T-ALL genome (**Table S1**). On a genomic organization perspective, these size estimates put hubs on a similar order of magnitude as TADs ^22^. To determine if T-ALL hubs were simply a subset of TADs, we identified TAD boundaries from DND41 Hi-C data and evaluated their overlap with hub boundaries. This analysis revealed that hubs were distinct from TADs with only 5.3% of the 2,754 hub boundaries overlapping with the more strongly insulated TAD boundaries (**Figs. S1a** and **S1b**), a conclusion that was corroborated by analysis of cohesin subunit SMC1 HiChIP (**Fig. S1c**). Given that hubs appeared to be unique from TADs, we characterized them by the count of spatial interactions between their constituent enhancer and promoter elements (**Fig. 1a**), where an interaction represents the presence of a contact between two regulatory elements (see Methods). As discussed in subsequent sections, we then leveraged Hi-C or HiChIP-measured enhancer and promoter interactivity within hubs, instead of individual algorithmically defined enhancer-promoter loops, as the basis for identifying differential hubs between two conditions (**Fig. 1b**).

To elucidate the basic organizational principles of enhancer-promoter hubs, we first examined within enhancer-promoter hub interaction counts. This analysis revealed that T-ALL hubs identified from either Hi-C or SMC1 HiChIP distributed asymmetrically, with only a small number of hubs harboring substantial numbers of spatial interactions (**Figs. 2a** and **S1d**). While 50.5% of Hi-C enhancer-promoter hubs contained less than 20 interactions, 11.5% or 158 hubs demonstrated high interactivity with more than 83 interactions in T-ALL (**Table S1**). Inspection of the relationship between number of promoters and enhancers participating in each hub and the extent of hub interactivity indicated that there was a positive correlation between interaction and regulatory element counts (**Figs. 2b** and **S1e**) such that hubs with the most interactions also tended to have the highest interaction to promoter/enhancer ratios (**Figs. 2c** and **S1f**). These observations suggest that the largest T-ALL hubs contain regulatory elements that are highly interacting, leading us to term this subset of enhancer-promoter assemblies as *hyperinteracting* hubs and refer to non-hyperinteracting hubs as *regular* hubs.

**Figure 2:**
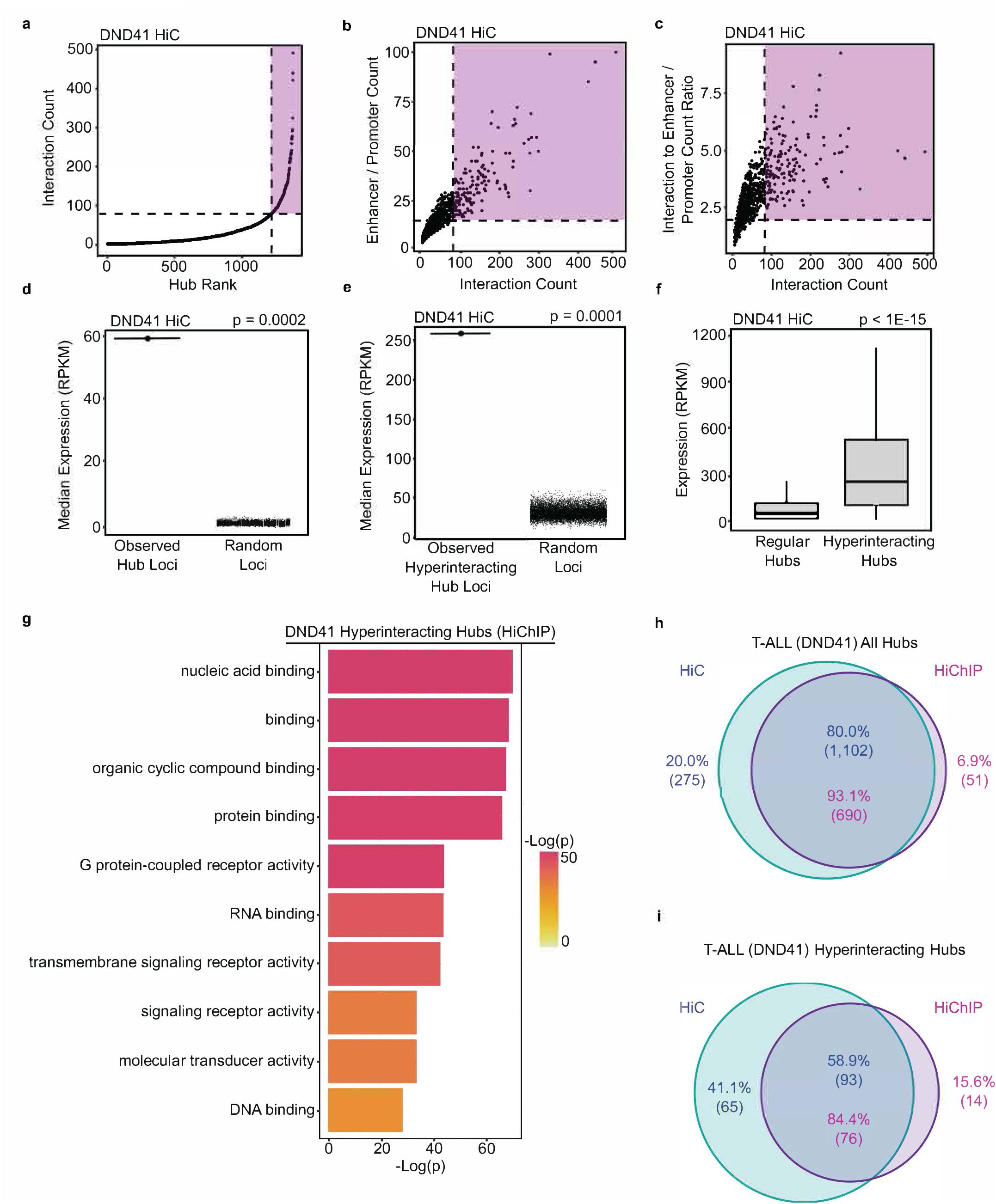
T-ALL hyperinteracting hubs are markedly transcribed and organize expression of genes encoding transcription factors and cofactors. a: Enhancer-promoter hubs detected from T-ALL DND41 Hi-C data are plotted in ascending order of their total interactivity. Ranking of enhancer-promoter hubs’ total interactivity reveals asymmetrical distribution of interactions among enhancers and promoters across the T-ALL genome, and delineates two classes of regular and hyperinteracting hubs. Hyperinteracting hubs are marked within the purple region and defined as the hubs above the elbow of the interactivity curve. b: Plotting Hi-C interaction count vs. enhancer/promoter element count in each T-ALL DND41 hub demonstrates that the number of interactions increases concordantly with the number of enhancer/promoter elements in enhancer-promoter hubs. The purple region marks hyperinteracting hubs. c: Plotting Hi-C interaction count vs. ratio of Hi-C interaction to enhancer/promoter counts shows that the T-ALL DND41 enhancer-promoter hubs with more interactions have more interconnected enhancer-promoter nodes, further supporting their hyperinteractivity. The purple region marks hyperinteracting hubs. d: T-ALL DND41 hubs are markedly enriched for transcriptional activity. Median expression of genes within loci of DND41 Hi-C hubs is compared with a permutation model as indicated by black points. Each black point corresponds to median gene expression observed in an iteration of the permutation model, which is comprised of 5,000 sets of randomly selected genomic regions with chromosomes, genomic lengths, and loci counts matching those of the DND41 hubs. P-value: empirical p-value. e: T-ALL DND41 hyperinteracting hubs are markedly enriched for transcriptional activity. Median expression of genes within loci of DND41 Hi-C hyperinteracting hubs is compared with a permutation model as indicated by black points. Each black point corresponds to median gene expression observed in an iteration of the permutation model, which is comprised of 10,000 sets of randomly selected genomic regions with chromosomes, genomic lengths, and loci counts matching those of the DND41 hyperinteracting hubs. P-value: empirical p-value. f: Box-and-whisker plots comparing transcription levels of T-ALL DND41 Hi-C regular and hyperinteracting hubs show that hyperinteracting hubs are significantly more transcribed than regular hubs (without genomic span normalization). Box-and-whisker plots: center line, median; box limits, upper (75th) and lower (25th) percentiles; whiskers, 1.53 interquartile range. P-value: two-tailed Wilcoxon rank sum test. g: Gene ontology (GO) enrichment analysis showing that activities characteristic of transcription factors and cofactors are among the top 10 most enriched molecular functions associated with expressed genes located in T-ALL DND41 SMC1 HiChIP hyperinteracting hubs. h: Venn diagram illustrating overlap of T-ALL DND41 Hi-C and SMC1 HiChIP hubs showing that Hi-C data is adequate to identify almost all of the hubs (93.1%) identified from higher resolution SMC1 HiChIP. Percentages and counts of overlapping and unique hubs are shown separately for Hi-C and SMC1 HiChIP. i: Venn diagram illustrating overlap of T-ALL DND41 Hi-C and SMC1 HiChIP hyperinteracting hubs showing that Hi-C data is adequate to identify most of the hyperinteracting hubs (84.4%) identified from higher resolution SMC1 HiChIP. Percentages and counts of overlapping and unique hyperinteracting hubs are shown separately for Hi-C and SMC1 HiChIP.

Given that both regular and hyperinteracting hubs are defined by the presence of active enhancers, we considered the possibility that they could be akin to super-enhancers. To examine the relationship between super-enhancers and enhancer-promoter hubs, we compared their location on the linear genome, observing that 60.5% of regular hubs did not coincide with super-enhancers. Importantly, 71.1% of hyperinteracting hubs contained two or more super-enhancers (**Fig. S1g**). This data indicates that spatial hubs are distinct from super-enhancers and suggests the potential enhanced regulatory status of hyperinteracting hubs in comparison to regular hubs.

### Hyperinteracting hubs organize control of transcription factor expression in T leukemic cells

Postulating that enhancer-promoter hubs function to tightly direct transcription of certain genes, we sought to examine the transcriptional status of T-ALL hubs. To this end, we evaluated RNA enrichment over hub loci in comparison to random representative loci of identical genomic length using total transcript RNA-seq in DND41 T-ALL cells. Our analysis showed that median RNA enrichment over the observed hubs was significantly greater than median enrichment in all of the 5,000 sets of size-matched random representative loci that served as comparators for Hi-C and SMC1 HiChIP hubs (**Fig. 2d**, p = 0.0002; and **Fig. S1h**, p = 0.0002). Together, these data suggest that T-ALL hubs are uniquely associated with transcriptional activation.

The observation that enhancer-promoter hubs are enriched for transcription in T-ALL led us to test if hyperinteracting hubs organize transcription on a broader scale than regular hubs and whether these two hub types are functionally distinct. Compared to regular hubs, SMC1 HiChIP hyperinteracting hubs on average spanned ∼3.2 times more base pairs (**Fig. S1i,** p = 1E-53). Examining transcription over hyperinteracting hubs revealed that median expression over these hubs was significantly higher than median expression over 10,000 sets of size-matched random representative loci from across the DND41 T-ALL genome (**Fig. 2e**, p = 0.0001; and **Fig. S1j**, p = 0.0001). Given that both regular and hyperinteracting hubs were distinctly enriched for transcription in comparison to random representative loci (**Figs. 2d**, **2e**, **S1h**, and **S1j**), we aimed to clarify their transcriptional state relative to one another. Consequently, we observed that hyperinteracting hubs contained more highly expressed genes (**Fig. S1k**, p = 1E-25) and were significantly enriched for gene expression (**Fig. 2f**, p < 1E-15) compared to regular hubs detected from Hi-C data, an observation that was corroborated by SMC1 HiChIP-measured interactions among enhancers and promoters (**Fig. S1l**, p < 1E-15). These findings support that hyperinteracting hubs coalesce enhancers and promoters to potentially establish transcriptionally permissive environments.

In light of these observations, we sought to determine the molecular functions of the genes contained within these highly transcribed T-ALL hyperinteracting hubs. Gene ontology (GO) enrichment analysis revealed that a significant fraction of expressed genes located within these topological assemblies encoded proteins functioning as transcription factors and cofactors (**Fig. 2g**; p < 1E-20; and **Table S2**), including DNA binding factors *MYC*, *TP53*, and *YY1* as well as chromatin and transcriptional coregulators *HDAC4*, *KAT5*, *DNMT3B*, and *EZH1* (**Table S1**) with demonstrated role in leukemia ^23, 24, 25, 26, 27^. As expected from the high degree of concordance between Hi-C and SMC1 HiChIP hubs (**Figs. 2h** and **2i**), GO enrichment analysis with hyperinteracting hubs identified from Hi-C corroborated these hubs’ distinct enrichment for transcription factors (**Table S2**). Collectively, these data suggest that hyperinteracting hubs may serve to not only organize local transcription, but also control transcription across the genome by regulating transcription factors that act in *trans* upon distal loci.

To assess whether our observations in DND41 were generalizable to other T-ALL models, we performed hub analysis in CUTLL1 cells ^28, 29, 30^. Similar to DND41, we observed asymmetrical distribution of interactions among enhancers and promoters and formation of hyperinteracting hubs in CUTLL1 (**Figs. S2a-c**). Moreover, CUTLL1 hubs were distinct from super-enhancers (**Fig. S2d**) and significantly transcribed (**Fig. S2e**, p = 0.0002; and **Fig. S2f**, p = 0.0001) with hyperinteracting hubs distinctly enriched for gene expression (**Fig. S2g**, p < 1E-15) and encoding molecules involved in nucleic acid / protein binding and receptor signaling (**Fig. S2h**, p < 1E-20).

To better understand the potential regulatory environment formed by hyperinteracting hubs in T-ALL, we next examined these hubs at histone methyltransferase *DOT1L* and DNA methyltransferase *DNMT3B* (**Figs. S2i** and **S2j**), two well-known genes involved in leukemogenesis with prognostic significance ^24, 31, 32^. Both *DOT1L* and *DNMT3B* hyperinteracting hubs contained several long-range interactions among highly accessible genomic elements with elevated levels of active histone mark H3K27ac and/or transcription. The *DOT1L* locus was among the top 10 most interacting hubs in both DND41 and CUTLL1 and exhibited similar local interaction organization. On the other hand, in the hyperinteracting hub containing *DNMT3B*, a number of active elements adjacent to the *DNMT3B* promoter demonstrated more variable spatial interactivity between the two T-ALL cell lines (**Figs. S2i** and **S2j**). Taken in conjunction, these data support the ability of hub-based analysis to detect key topological assemblies with varying spatial interaction structure across different models of a given cancer.

### Organizational principles of enhancer-promoter hubs are shared between T cell leukemia and B cell lymphoma

Observing organizational principles of highly interacting enhancer-promoter hubs in T-ALL led us to investigate whether these topological assemblies create complex networks of regulatory elements interacting with genes encoding transcriptional regulators in other cancer types. For this reason, we identified hubs from Rec-1 MCL cells using both Hi-C and SMC1 HiChIP data ^14^. Similar to T-ALL, Rec-1 hubs were largely distinct from both TADs and super-enhancers (**Figs. S3a-d**), and demonstrated significant asymmetry in interactivity and enhancer/promoter count distributions (**Figs. 3a, 3b, S3e**, and **S3f**). The existence of a positive correlation between interaction counts and interaction to enhancer/promoter ratios further supports the presence of hyperinteracting hubs in MCL (**Figs. 3c** and **S3g**). Rec-1 hubs in general (**Fig. 3d**, p = 0.0002; and **Fig. S3h**, p = 0.0002) and hyperinteracting hubs in particular (**Fig. 3e**, p = 0.0001; and **Fig. S3i**, p = 0.0001) both demonstrated significant transcriptional activity. In comparison to regular hubs, Rec-1 hyperinteracting hubs spanned significantly more base pairs on the linear genome (**Fig. S3j**, p = 1E-31), contained increased numbers of highly expressed genes (**Fig. S3k**, p = 1E-32), and exhibited markedly more transcriptional activity than regular hubs (**Fig. 3f**, p < 1E-15; and **Fig. S3l**, p < 1E-15), similar to T-ALL.

**Figure 3:**
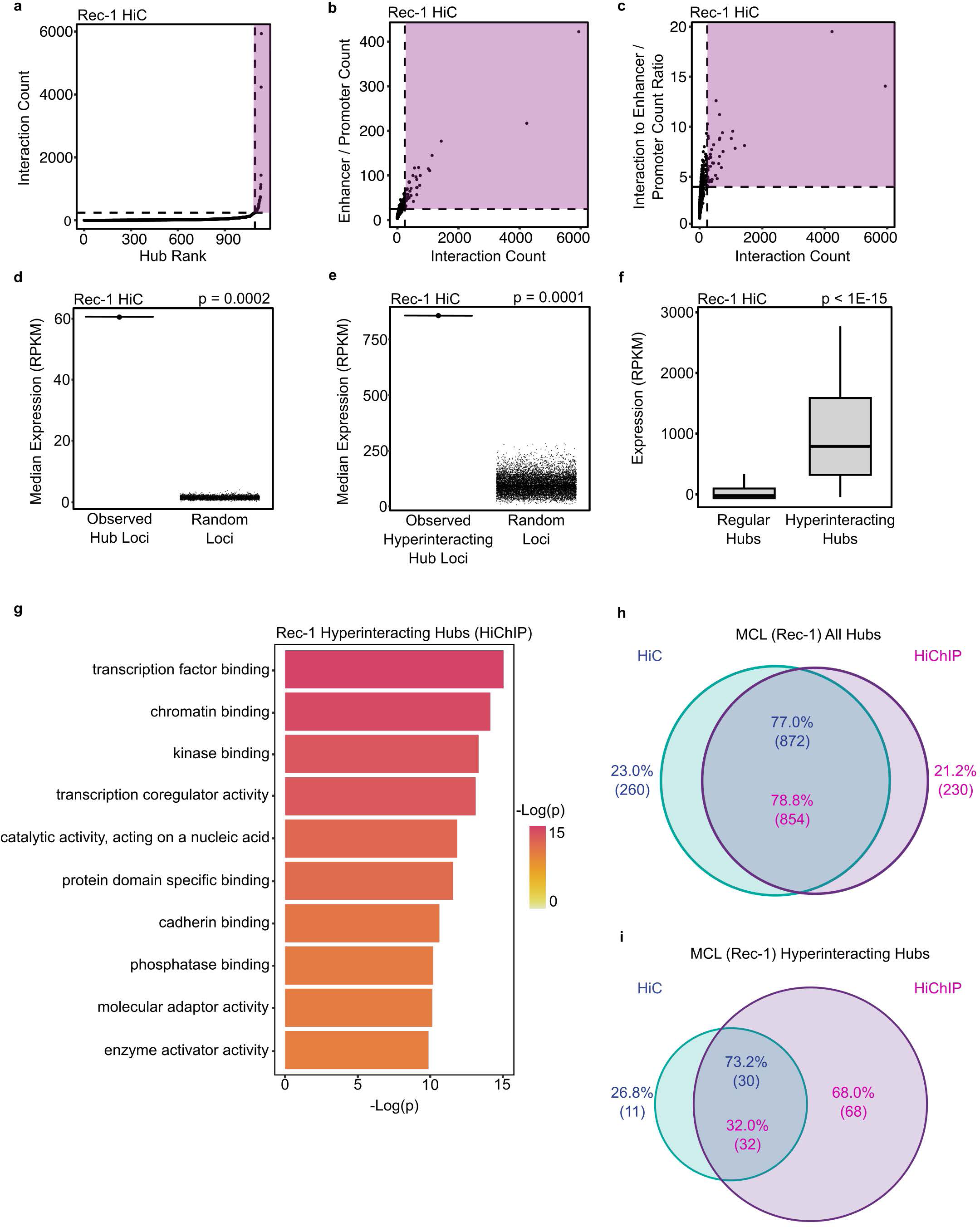
MCL hyperinteracting hubs are markedly transcribed and organize expression of genes encoding transcription factors and cofactors. a: Enhancer-promoter hubs detected from MCL Rec-1 Hi-C data are plotted in ascending order of their total interactivity. Ranking of enhancer-promoter hubs’ total interactivity reveals asymmetrical distribution of interactions among enhancers and promoters across the MCL genome, and delineates two classes of regular and hyperinteracting hubs. Hyperinteracting hubs are marked within the purple region and defined as the hubs above the elbow of the interactivity curve. b: Plotting Hi-C interaction count vs. enhancer/promoter element count in each MCL Rec-1 hub demonstrates that the number of interactions increases concordantly with the number of enhancer/promoter elements in enhancer-promoter hubs. The purple region marks hyperinteracting hubs. c: Plotting Hi-C interaction count vs. ratio of Hi-C interaction to enhancer/promoter counts shows that the MCL Rec-1 enhancer-promoter hubs with more interactions have more interconnected enhancer-promoter nodes, further supporting their hyperinteractivity. The purple region marks hyperinteracting hubs. d: MCL Rec-1 hubs are markedly enriched for transcriptional activity. Median expression of genes within loci of Rec-1 Hi-C hubs is compared with a permutation model as indicated by black points. Each black point corresponds to median gene expression observed in an iteration of the permutation model, which is comprised of 5,000 sets of randomly selected genomic regions with chromosomes, genomic lengths, and loci counts matching those of the Rec-1 hubs. P-value: empirical p-value. e: MCL Rec-1 hyperinteracting hubs are markedly enriched for transcriptional activity. Median expression of genes within loci of Rec-1 Hi-C hyperinteracting hubs is compared with a permutation model as indicated by black points. Each black point corresponds to median gene expression observed in an iteration of the permutation model, which is comprised of 10,000 sets of randomly selected genomic regions with chromosomes, genomic lengths, and loci counts matching those of the Rec-1 hyperinteracting hubs. P-value: empirical p-value. f: Box-and-whisker plots comparing transcription levels of MCL Rec-1 Hi-C regular and hyperinteracting hubs show that hyperinteracting hubs are significantly more transcribed than regular hubs (without genomic span normalization). Box-and-whisker plots: center line, median; box limits, upper (75th) and lower (25th) percentiles; whiskers, 1.53 interquartile range. P-value: two-tailed Wilcoxon rank sum test. g: Gene ontology (GO) enrichment analysis showing that activities characteristic of transcription factors and cofactors are among the top 10 most enriched molecular functions associated with expressed genes located in MCL Rec-1 SMC1 HiChIP hyperinteracting hubs. h: Venn diagram of overlap of MCL Rec-1 Hi-C and SMC1 HiChIP hubs showing that Hi-C data is adequate to identify most of the hubs (78.8%) identified from higher resolution SMC1 HiChIP. Percentages and counts of overlapping and unique hubs are shown separately for Hi-C and SMC1 HiChIP. i: Venn diagram of overlap of MCL Rec-1 Hi-C and SMC1 HiChIP hyperinteracting hubs showing that Hi-C data identifies 32.0% of the hyperinteracting hubs identified from higher resolution SMC1 HiChIP. This difference is likely due to the presence of two outlier hubs from Hi-C data, which leads to a more stringent cutoff for hyperinteracting hubs in Hi-C compared to SMC1 HiChIP data. Percentages and counts of overlapping and unique hyperinteracting hubs are shown separately for Hi-C and SMC1 HiChIP.

To examine the potential role of genes located in MCL hyperinteracting hubs, we performed molecular function GO enrichment analysis. This analysis revealed that, in agreement with T-ALL, a significant fraction of genes located within MCL hyperinteracting hubs encoded transcription factors and cofactors (**Fig. 3g**, p < 1E-9; and **Table S2**), including *MYC*, *CTCF*, *ETS1*, *KAT5*, and *DOT1L*. These genes were located in hyperinteracting hubs in both Rec-1 MCL and DND41 T-ALL cells (**Table S1**); yet, the structure of these hubs appeared different between the two cancer types, as exemplified by the *DOT1L* locus (**Fig. S3m**). On the other hand, several transcription regulators with roles in oncogenesis, such as *DNMT3B* ^24^ (**Figs. S3n** and **S3o**; **Table S1**) and B lymphocyte lineage transcription factor *PAX5* ^33^ (**Figs. S3p** and **S3q**; **Table S1**) were only expressed and positioned at hyperinteracting hubs in either T-ALL or MCL and not both. As such, these data corroborate observations in T-ALL (**Fig. 2g**) and further support the role of hyperinteracting hubs as potential regulatory assemblies that orchestrate gene expression in hematological malignancies.

### Enhancer-promoter hub identification is robust to chromatin conformation capture assay resolution

*In-situ* Hi-C provides unbiased chromatin conformation maps at the expense of short-range enhancer-promoter loop resolution. In contrast, protein-centric assays, including HiChIP, are biased to increase resolution and support identification of looping interactions mediated by a particular protein ^34^. To assess the impact of chromatin conformation capture technology on detection of enhancer-promoter hubs, we first compared hubs identified with Hi-C and SMC1 HiChIP in DND41 T-ALL. Hubs detected with SMC1 HiChIP in DND41 were fewer and on average larger than hubs detected with Hi-C. 741 SMC1 HiChIP hubs were identified with an average span of 1.09 Mb compared to 1,377 Hi-C hubs spanning an average of 414 Kb. A similar trend was observed for hyperinteracting hubs, where SMC1 HiChIP and Hi-C identified 90 and 158 hyperinteracting hubs with an average span of 2.73 Mb and 1.00 Mb on the linear DND41 genome, respectively (**Table S1**). Despite these differences, the vast majority of hubs identified with SMC1 HiChIP were also detectable with Hi-C, where 93.1% of total SMC1 HiChIP hubs and 84.4% of hyperinteracting SMC1 HiChIP hubs specifically coincided with Hi-C total and hyperinteracting hubs, respectively (**Figs. 2h** and **2i**). Hence, DND41 SMC1 HiChIP hyperinteracting hubs were largely a subset of Hi-C hubs. While 84.4% of SMC1 HiChIP hyperinteracting hubs overlapped with Hi-C hyperinteracting hubs, only 58.9% of Hi-C hyperinteracting hubs overlapped with their SMC1 HiChIP counterparts (**Fig. 2i**). As shown in Figures 2a to 2f and S1d-f, S1h, S1j, and S1l, both hubs detected from SMC1 HiChIP and Hi-C exhibited similar transcriptional activity as well as interaction and enhancer/promoter count distributions. In sum, these data exhibit the high fidelity of our analysis to identify enhancer-promoter hubs in T-ALL from both Hi-C and SMC1 HiChIP experiments.

In order to corroborate these observations, we repeated comparative analyses with SMC1 HiChIP and Hi-C data from Rec-1 MCL cells. Similar to DND41 T-ALL, Hi-C hubs coincided with more than 75% of HiChIP hubs (**Fig. 3h**). However, the percentage of Rec-1 SMC1 HiChIP hyperinteracting hubs overlapping with Hi-C hyperinteracting hubs was only 32.0% (**Fig. 3i**), which was likely due to two Hi-C outlier hubs with disproportionately large interaction counts (**Figs. 3a** and **S3r**) that resulted in a more stringent cutoff for categorizing a hub as hyperinteracting from Hi-C compared to HiChIP measurements. Nevertheless, Rec-1 Hi-C and SMC1 HiChIP hyperinteracting hubs exhibited similar structural and transcriptional characteristics (**Figs. 3a-f**, **S3e-i**, and **S3l**). In tandem with the results of our comparative analyses in T-ALL, these data suggest that hub analysis is robust across a spectrum of chromatin conformation capture assays with varying resolution of enhancer-promoter looping. More importantly, our analysis supports the role of enhancer-promoter hubs as potentially important units of genome organization rather than artifacts of a particular chromatin capture assay.

### Organizational principles of enhancer-promoter hubs in hematological cancers are generalizable to non-hematological cancers

Intrigued by the commonality of hub organizational principles in leukemia and lymphoma, we sought to evaluate structural and transcriptional characteristics of enhancer-promoter hubs in non-hematological cancers. To this end, we identified hubs in two triple-negative breast cancer (TNBC) cell lines, HCC1599 and MB157, using SMC1 HiChIP data ^14^. Analysis of TNBC enhancer-promoter hubs confirmed that they could be stratified on the basis of interaction count into two distinct groups of regular and hyperinteracting hubs (**Figs. 4a-c** and **S4a-c**), both of which were markedly transcribed (**Fig. 4d**, p = 0.0002; **Fig. 4e**, p = 0.0001; **Fig. S4d,** p = 0.0002; and **Fig. S4e**, p = 0.0001) and distinct from super-enhancers (**Figs. S4f** and **S4g**). Similar to T-ALL and MCL, hyperinteracting hubs in TNBC contained a greater number of highly expressed genes (**Fig. S4h**, p = 1E-23; and **Fig. S4i**, p = 1E-32). Furthermore, TNBC hyperinteracting hubs were significantly more transcribed (**Fig. 4f**, p = 1E-11; and **Fig. S4j**, p < 1E-15) and spanned larger genomic distances than regular hubs (**Fig. S4k**, p = 1E-23; and **Fig. S4l**, p = 1E-14), again mirroring T-ALL and MCL hubs. TNBC hyperinteracting hubs also predominantly formed at genes encoding transcription factors and cofactors (**Figs. 4g** and **S4m**; **Table S2**), some of which were only present in TNBC. For example, TNBC-associated hyperinteracting hubs formed around *SOX9* (**Figs. 4h** and **4i**), a transcription factor with demonstrated role in TNBC oncogenesis ^35, 36, 37^, and *TRPS1* (**Figs. S4n** and **S4o**), a transcription factor that serves as a highly specific marker for breast carcinoma including TNBC ^38, 39^. In contrast, some of the most prominent hyperinteracting hubs in T-ALL and/or MCL, including *DOT1L*, *DNMT3B*, and *PAX5* **(Figs. S3m, S3n**, and **S3p),** were not hyperinteracting or not present in TNBC (**Figs. S4p-r).** Taken together, our characterization of TNBC hubs corroborates T-ALL and MCL observations, and suggests that hyperinteracting hubs may organize transcriptional regulation of trans-acting factors in a lineage-associated manner to inform broader gene expression programs.

**Figure 4:**
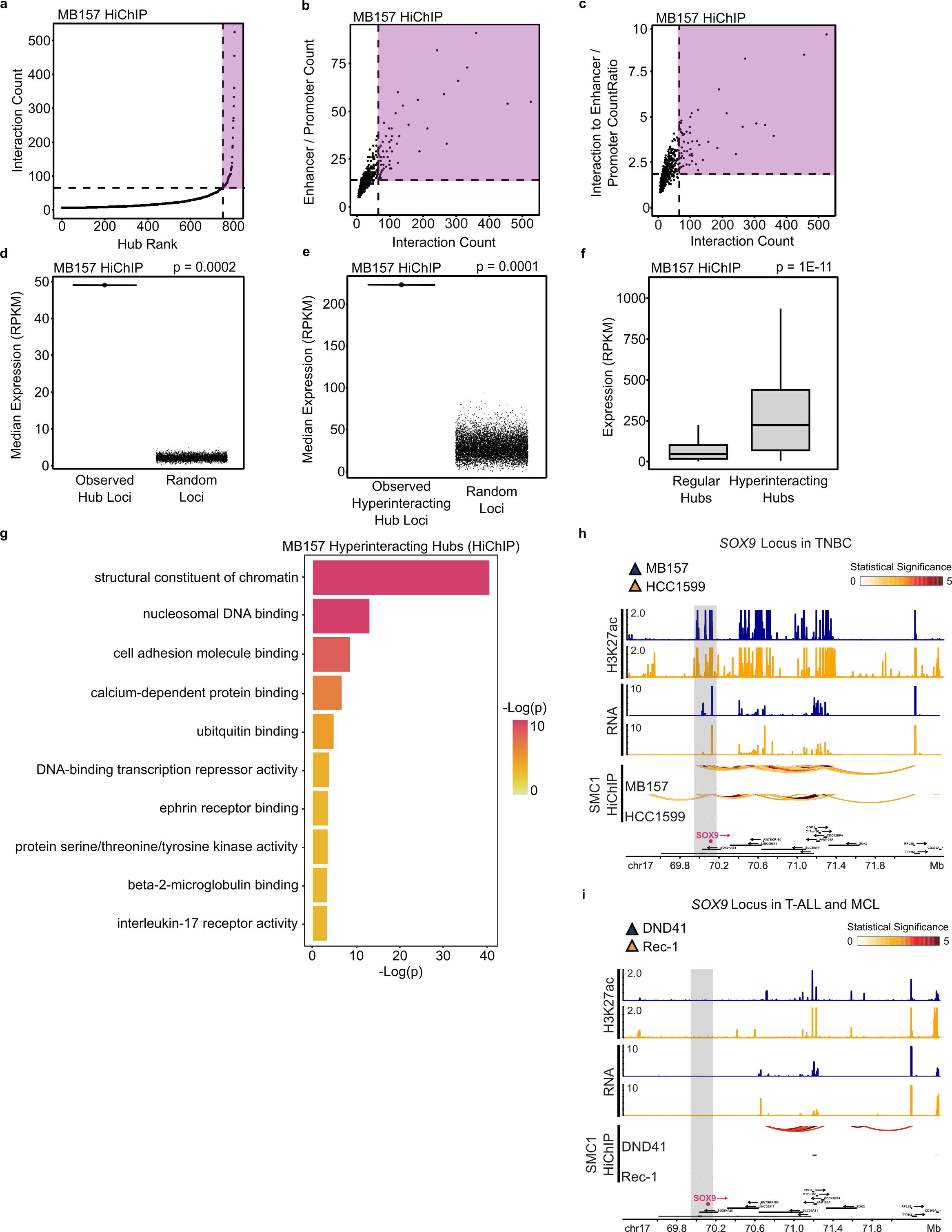
TNBC hyperinteracting hubs are markedly transcribed and organize expression of genes encoding transcription factors and cofactors. a: Enhancer-promoter hubs detected from TNBC MB157 SMC1 HiChIP data are plotted in ascending order of their total interactivity. Ranking of enhancer-promoter hubs’ total interactivity reveals asymmetrical distribution of interactions among enhancers and promoters across the TNBC genome, and delineates two classes of regular and hyperinteracting hubs. Hyperinteracting hubs are marked within the purple region and defined as the hubs above the elbow of the total interactivity ranking. b: Plotting SMC1 HiChIP interaction count vs. enhancer/promoter element count in each TNBC MB157 hub demonstrates that the number of interactions increases concordantly with the number of enhancer/promoter elements in enhancer-promoter hubs. The purple region marks hyperinteracting hubs. c: Plotting SMC1 HiChIP interaction count vs. ratio of SMC1 HiChIP interaction to enhancer/promoter counts shows that the TNBC MB157 enhancer-promoter hubs with more interactions have more interconnected enhancer-promoter nodes, further supporting their hyperinteractivity. The purple region marks hyperinteracting hubs. d: TNBC MB157 hubs are markedly enriched for transcriptional activity. Median expression of genes within loci of MB157 SMC1 HiChIP hubs is compared with a permutation model as indicated by black points. Each black point corresponds to median gene expression observed in an iteration of the permutation model, which is comprised of 5,000 sets of randomly selected genomic regions with chromosomes, genomic lengths, and loci counts matching those of the MB157 hubs. P-value: empirical p-value. e: TNBC MB157 hyperinteracting hubs are markedly enriched for transcriptional activity. Median expression of genes within loci of MB157 SMC1 HiChIP hyperinteracting hubs is compared with a permutation model as indicated by black points. Each black point corresponds to median gene expression observed in an iteration of the permutation model, which is comprised of 10,000 sets of randomly selected genomic regions with chromosomes, genomic lengths, and loci counts matching those of the MB157 hyperinteracting hubs. P-value: empirical p-value. f: Box-and-whisker plots comparing transcription levels of TNBC MB157 SMC1 HiChIP regular and hyperinteracting hubs show that hyperinteracting hubs are significantly more transcribed than regular hubs (without genomic span normalization). Box-and-whisker plots: center line, median; box limits, upper (75th) and lower (25th) percentiles; whiskers, 1.53 interquartile range. P-value: two-tailed Wilcoxon rank sum test. g: Gene ontology (GO) enrichment analysis showing that activities characteristic of transcription factors and cofactors are among the top 10 most enriched molecular functions associated with expressed genes located in TNBC MB157 SMC1 HiChIP hyperinteracting hubs. h, i: *SOX9* is located in a hyperinteracting hub in TNBC but not T-ALL and MCL. SMC1 HiChIP arcs show that the gray box-marked *SOX9* forms a hyperinteracting enhancer-promoter hub with several active regulatory elements and genes marked with H3K27ac ChIP-seq and RNA-seq, respectively, in TNBC MB157 and HCC1599 (h) but not T-ALL DND41 and MCL Rec-1 (i). Bottom tracks indicate Ensembl gene position.

### Hyperinteracting hubs are more lineage associated than regular hubs

The lineage association of certain key hyperinteracting hubs (**Figs. S2i-j**, **S3m-q**, **S4n-r,** and **4h-i**) led us to systematically assess the similarity of hubs identified from T-ALL, MCL, and TNBC SMC1 HiChIP data. By defining hub similarity as the percentage of hubs with overlapping genomic loci between two cancer types, we observed that hubs are relatively conserved, with at least 51% of hubs being shared between any two cancer types (**Fig. 5a**), and at least 23% of hubs being shared across all four cancer types (data not shown). Some of the common hubs in T-ALL, MCL, and TNBC contained genes encoding key transcription and DNA replication regulators including *TET3*, *CTCF*, *KLF10*, *AKT1*, *E2F1/4/6*, *PARP* enzymes, polymerases, and *NFkB* proteins (**Table S1**).

**Figure 5:**
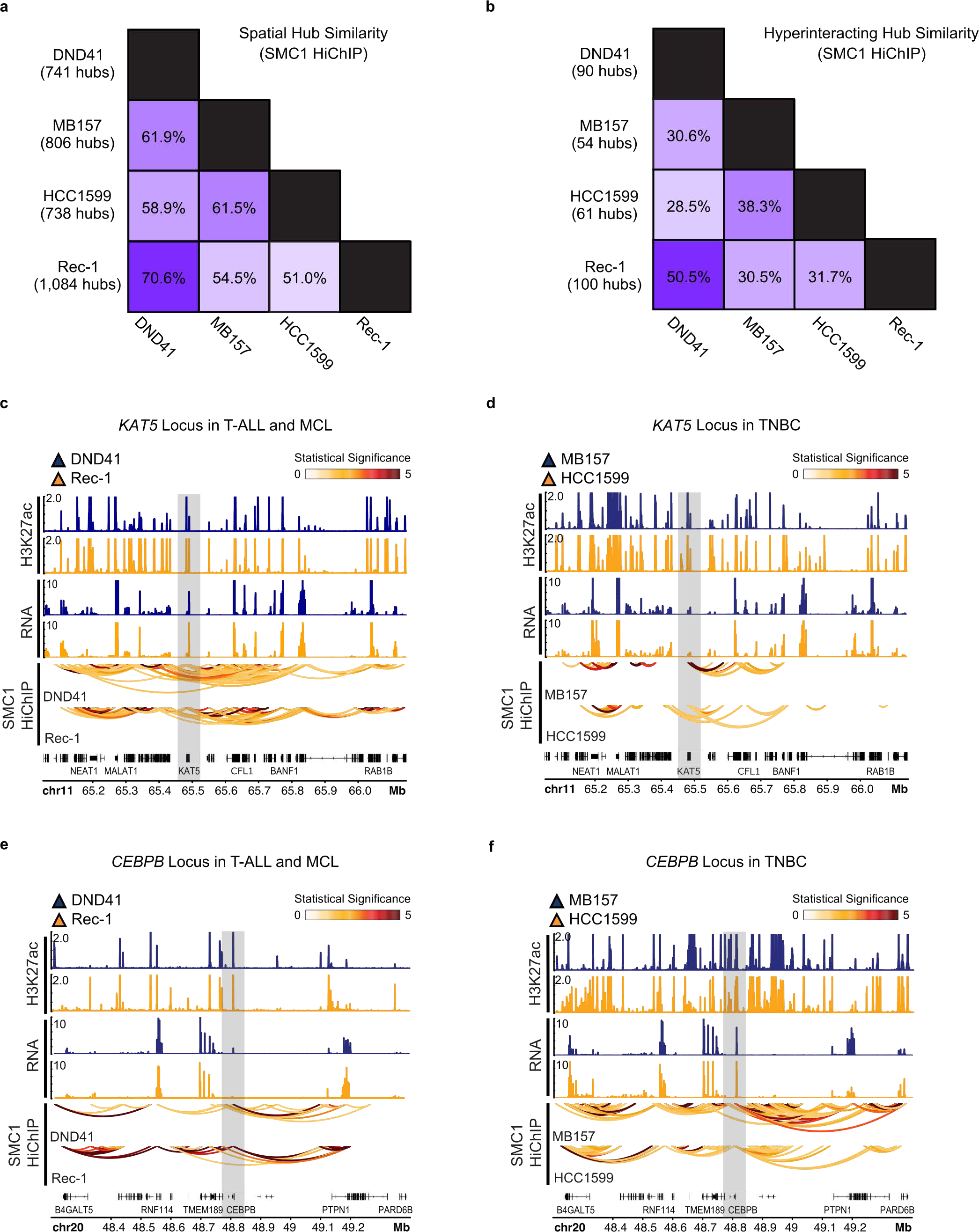
Hyperinteracting hubs are more lineage associated compared to regular hubs in T-ALL, MCL, and TNBC. a: Matrix of pairwise SMC1 HiChIP enhancer-promoter hub similarity shows that these topological assemblies are somewhat conserved across T-ALL, MCL, and TNBC cells. Similarity between each pair is calculated as the percentage of enhancer-promoter hubs with overlapping genomic intervals. Total hub counts for each cell line is listed on the left. b: Matrix of pairwise SMC1 HiChIP hyperinteracting hub similarity shows that these topological assemblies are more lineage associated than all enhancer-promoter hubs in T-ALL, MCL, and TNBC cells. Similarity between each pair is calculated as the percentage of hyperinteracting hubs with overlapping genomic intervals. Total hyperinteracting hub counts for each cell line is listed on the left. c, d: *KAT5* is located in a hyperinteracting hub in T-ALL and MCL but not TNBC. SMC1 HiChIP arcs show that the gray box-marked *KAT5* forms a hyperinteracting enhancer-promoter hub with several active regulatory elements and genes marked with H3K27ac ChIP-seq and RNA-seq, respectively, in T-ALL DND41 and MCL Rec-1 (c) but not TNBC MB157 and HCC1599 (d). Bottom tracks indicate Ensembl gene position. e, f: *CEBPB* is located in a hyperinteracting hub in TNBC but not T-ALL and MCL. SMC1 HiChIP arcs show that the gray box-marked *CEBPB* forms a hyperinteracting enhancer-promoter hub with several active regulatory elements and genes marked with H3K27ac ChIP-seq and RNA-seq, respectively, in TNBC MB157 and HCC1599 (f) but not T-ALL DND41 and MCL Rec-1 (e). Bottom tracks indicate Ensembl gene position.

Observing that hubs are generally shared across T-ALL, MCL, and TNBC, we went on to compare hyperinteracting hubs in these cancers. In contrast to all hubs, hyperinteracting hubs were more lineage associated such that less than ∼50% of these hubs were shared between any two cancer types (**Fig. 5b**). The observation that hyperinteracting hubs were more lineage associated than regular hubs was further supported by comparison of hubs detected from Hi-C data in DND41 and CUTLL1 T-ALL as well as Rec-1 MCL (**Figs. S5a** and **S5b**).

To evaluate lineage correlation of hyperinteracting hubs in greater depth, we closely scrutinized their organization across T-ALL, MCL, and TNBC. Lineage-associated hyperinteracting hubs were separated into two groups based on their presence and interactivity. The first group consisted of hyperinteracting hubs that only existed in one cancer type, as exemplified by the *SOX9, PAX5*, and *DNMT3B* genes, which were only coalesced within enhancer-promoter hubs in TNBC, MCL, and T-ALL, respectively (**Figs. 4h**, **4i**, S3n, S3p, and **S4q-r**). The second group consisted of hubs that were highly interacting in some cancer types and less interacting in others, as exemplified by the *DOT1L*, *KAT5* (also known as *TIP60*), and *CEBPB* genes. Similar to the hub containing *DOT1L* (**Figs. S3m** and **S4p**), the hub containing *KAT5*, a histone acetyltransferase with known role in driving *HOXA* gene expression in leukemia ^23, 40^, was hyperinteracting in T cell leukemia and B cell lymphoma but regularly interacting in TNBC (**Figs. 5c** and **5d**). In contrast, the hub containing *CEBPB*, a transcription factor implicated in normal and malignant breast epithelium ^41^, was regularly interacting in leukemia and lymphoma, but was hyperinteracting in TNBC (**Figs. 5e** and **5f**). Other notable hyperinteracting hub genes that were spatially proximate to multiple regulatory elements in T-ALL and MCL but to only a few in TNBC included the retinoic acid receptor *RXRB*, the apoptosis regulator *BAK1*, and the heat shock protein *HSPA9* (also known as mortalin; **Table S1**). On the other hand, as previously discussed, the transcription factor *TRPS1* was located within a hyperinteracting hub only in TNBC and not T-ALL or MCL (**Figs. S4n** and **S4o**). Together, these data suggest that hyperinteracting hubs are a distinct subset of enhancer-promoter hubs that demonstrate notable lineage association and are potentially involved in transcriptional control.

Despite observing that most hyperinteracting hubs were lineage associated, we identified ten hyperinteracting hubs that were present in T-ALL, MCL, and TNBC (**Fig. S5c**). Some of these topological assemblies, which were conserved on the basis of genomic overlap, formed at genes implicated in oncogenesis. Notable expressed genes from these hubs include the proto-oncogene *MYC*, the *MYC*-interacting chromatin effector *PYGO2* ^42^, the cell proliferation driver *EFNA4* ^43, 44^, the well-studied protein deacetylase *SIRT2* ^45^, and the highly expressed transcription factor and cancer biomarker *ZNF217* ^46^ (**Figs. S5c-e**). The intriguing commonality of these few hyperinteracting hubs highlights the potential importance of enhancer-promoter hubs in oncogenesis.

### Enhancer-promoter hubs are reorganized in GSI-resistant T-ALL

Given the correlation between enhancer-promoter hubs’ interactivity and transcriptional level in T-ALL, MCL, and TNBC, we aimed to examine the potential structure-function relationship of hubs by studying their changes during anticancer drug resistance acquisition. To this end, we first screened for differential hubs in T leukemic cells that were either sensitive or resistant to gamma secretase inhibitor (GSI), an antagonist of NOTCH1 signaling, with *NOTCH1* being the most frequently mutated gene in T-ALL ^47, 48^. Differential hubs were identified based on within-hub interaction count changes between GSI-sensitive and GSI-resistant DND41 T-ALL cells (**Fig. 1b** and **Table S3**) (see Methods). We postulated that identifying differential hubs should reveal genes with key roles in GSI resistance without the need for differential loop calling analysis, should hubs function as topological assemblies of gene expression control.

Differential hub analysis in GSI-sensitive and GSI-resistant DND41 T-ALL identified 217 differential hubs that had at least 2-fold change in interactivity or were *de novo* gained/lost in GSI resistance (**Fig. 6a**). Notably, these differential hubs were distinct from differential compartments and TADs identified in

**Figure 6:**
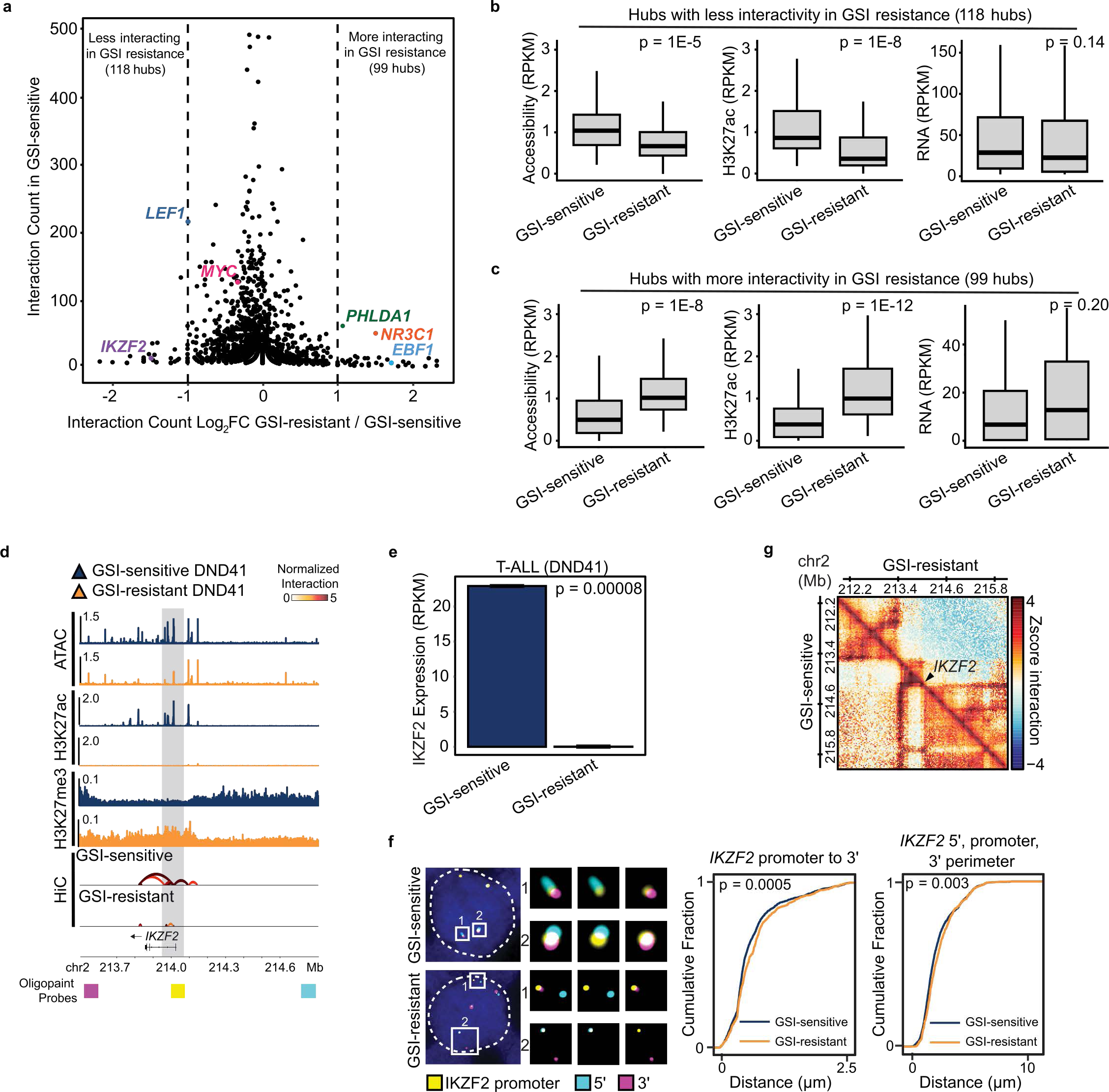
Loss of *IKZF2* hyperinteracting hub coincides with decrease in chromatin activity, gene expression, and architectural stripe in GSI-resistant T-ALL. a: Enhancer-promoter hubs with differential interactivity in GSI-resistant vs. -sensitive T-ALL DND41 contain genes with potential roles in oncogenesis and / or treatment response. Scatter plot showing log_2_ fold change of hub interactivity in GSI-resistant and GSI-sensitive DND41 (x axis) vs. hub interaction counts in GSI-sensitive cells. Dotted lines mark enhancer-promoter hubs with more than or equal to 2-fold decrease (“less interacting in GSI resistance”) or increase (“more interacting in GSI resistance”) in interaction counts in GSI-resistant vs. GSI-sensitive DND41 cells. Hubs that were *de novo* gained or lost in GSI resistance based on the hub minimum interactivity threshold of 6 were also considered differential. Selected hubs containing genes with potential role in oncogenesis and / or treatment response are labeled. For visual clarity, log_2_ fold changes with pseudocount 1 are shown. b: Box-and-whisker plots showing significant reduction in chromatin accessibility (left) and H3K27ac deposition (center), with a similar trend for gene expression (right) in hubs that lost interactivity in GSI-resistant cells. Box-and-whisker plots: center line, median; box limits, upper (75th) and lower (25th) percentiles; whiskers, 1.53 interquartile range. P-value: two-tailed Wilcoxon rank sum test. c: Box-and-whisker plots showing significant increase in chromatin accessibility (left) and H3K27ac deposition (center), with a similar trend for transcription (right) in hubs that gained interactivity in GSI-resistant cells. Box-and-whisker plots: center line, median; box limits, upper (75th) and lower (25th) percentiles; whiskers, 1.53 interquartile range. P-value: two-tailed Wilcoxon rank sum test. d: Genome tracks at the *IKZF2* locus show that loss of Hi-C interactions in the *IKZF2* hyperinteracting hub in GSI-resistant DND41 cells coincide with marked decrease in chromatin accessibility and H3K27ac active chromatin mark as well as gain of H3K27me3 heterochromatic mark. Note that the arcs displayed in this illustration represent only the valid interactions detected from Hi-C between enhancer and/or promoter elements (with normalized contact frequency > 3). Oligopaint DNA FISH probes are marked with pseudo-color magenta (*IKZF2* 3’), yellow (gray box-marked *IKZF2* promoter), and cyan (*IKZF2* 5’) below the Ensembl gene track. e: *IKZF2* is significantly downregulated in GSI-resistant T-ALL cells. Barplots of normalized RNA-seq reads showing *IKZF2* mRNA levels in GSI-resistant and GSI-sensitive cells. N: 3; P-value: unpaired student t-test; error bars: +/- 2 standard error of the mean (SE). f: *IKZF2* enhancer-promoter hub expands in GSI-resistant T-ALL DND41 cells. Left: confocal microscopy images of representative cells and magnified images for 3-color *IKZF2* 3’ (magenta), *IKZF2* promoter (yellow), *IKZF2* 5’ (cyan) probes in GSI-sensitive (top) and GSI-resistant (bottom) cells are shown. Locations of three 75 Kb Oligopaint probes *IKZF2* 3’ (magenta), *IKZF2* promoter (yellow), *IKZF2* 5’ (cyan) are marked in (D). Center: cumulative distribution plots of the closest distance between the *IKZF2* promoter and its 3’ locus in the same cell are compared between 1548 GSI-sensitive and 533 GSI-resistant allelic interactions. Mean (+/- S.D.) distance in GSI-sensitive and GSI-resistant cells is 0.642 (+/- 0.50) µm, and 0.704 (+/- 0.52) µm, respectively (Kolmogorov-Smirnov p-value = 0.0005). Right: cumulative distribution plots of spatial perimeter formed by the *IKZF2* 3’, promoter, and 5’ probes. Mean (+/- S.D.) in GSI-sensitive and GSI-resistant cells is 2.26 (+/- 1.31) µm, and 2.41 (+/- 1.27) µm, respectively (Kolmogorov-Smirnov p-value = 0.0027). g: Normalized Hi-C contact maps in GSI-resistant (upper triangle) and GSI-sensitive cells (lower triangle) at the *IKZF2* locus shows the loss of an architectural stripe connected to *IKZF2*.

GSI-sensitive and -resistant cells (**Figs. S6a-d**) in accordance with enhancer-promoter hubs’ broader separation from TADs and super-enhancers (**Fig. S1a-c**, **S1g**, and **S3a-d**). To determine whether these spatially differential hubs were also epigenetically or transcriptionally altered in GSI resistance, we examined their differential chromatin accessibility, chromatin activity, and gene expression using ATAC-seq, H3K27ac ChIP-seq, and RNA-seq, respectively. Interestingly, hubs with significant loss or gain of interactivity in GSI resistance demonstrated markedly decreased or increased chromatin opening (**Fig. 6b**, p = 1E-5; and **Fig. 6c**, p = 1E-8) and chromatin activity (**Fig. 6b**, p = 1E-8; and **Fig. 6c**, p = 1E-12), respectively, with a similar trend for gene expression (**Fig. 6b**, p = 0.14; and **Fig. 6c**, p = 0.20). Absence of statistically significant changes in hubs’ transcriptional activity could be in part attributed to lack of concordant changes in all the genes located within these topological assemblies such that aggregate measurements of hub transcription appear less variant. Nonetheless, close examination of hubs with a gain of interactivity in GSI-resistant T leukemic cells showed that some of these topological assemblies formed at B cell-related genes, including Early B-cell factor 1 (*EBF1;* **Table S3**), as well as genes involved in glucocorticoid signaling, including glucocorticoid receptor *NR3C1* (**Table S2**). Transcription factor *EBF1*, which promotes development of GSI resistance in T-ALL ^21^, is both more interacting (**Fig. 6a**) and expressed (**Fig. S6e**, p = 0.003) in GSI-resistant compared to GSI-sensitive cells. Similarly, *NR3C1*, a key driver of T-ALL steroid resistance ^49^, participated in an enhancer-promoter hub with more than a 2.5-fold gain of interactivity (**Fig. 6a**) and expression (**Fig. S6f**, p = 0.003) in the GSI-resistant state. On the other hand, scrutinization of hubs with loss of interactivity in GSI-resistant cells revealed disruption of highly interacting enhancer-promoter assemblies at genes involved in T cell biology and T cell receptor signaling (**Table S2**), including *LEF1* and *IKZF2*. These data are supported by earlier findings showing that T-ALL GSI resistance is partially mediated by downregulation of a T cell-related transcription program in favor of a B cell-related one ^21^, hence reinforcing the potential functional importance of enhancer-promoter hub restructuring during GSI resistance.

### Optical mapping confirms reorganization of enhancer-promoter hubs at individual GSI-resistant T-ALL cells

Given that downregulation of a T cell-associated transcription program spurs acquisition of GSI resistance in T-ALL ^21^, we closely examined hubs at T cell lineage-restricted transcription factors *LEF1* and *IKZF2*, which experienced nearly 2- and 3-fold reductions in interactivity in GSI-resistant T-ALL, respectively (**Figs. S6g** and **6d, Table S3**). In line with decreased interactivity at these two loci, we observed marked reduction in chromatin accessibility, substantial depletion of active enhancer mark H3K27ac, and deposition of repressive mark H3K27me3 (**Figs. S6g** and **6d**), which was concomitant with significant repression of *LEF1* (**Fig. S6h**; p = 0.0003) and *IKZF2* (**Fig. 6e**; p = 0.00008).

We next sought to establish how reduction of interaction frequency at *LEF1* and *IKZF2* enhancer-promoter hubs detected from Hi-C relates to physical separation of regulatory elements within these two loci in individual GSI-resistant cells. To this end, we used high-resolution Oligopaint DNA fluorescence *in situ* hybridization (FISH) and 3D confocal imaging to visualize physical perimeters of three genomic elements at each locus in GSI-sensitive and GSI-resistant DND41 cells.

To detect the disruption of *LEF1* at a single-cell resolution, we designed Oligopaint DNA FISH probes hybridizing to three regulatory elements surrounding *LEF1*: the *LEF1* promoter (5’ end of the locus), a lineage-restricted *LEF1* enhancer (center of the locus), and the *RPL34* promoter (3’ end of the locus). Measuring pairwise distances between the *LEF1* promoter, the *RPL34* promoter, and the lineage-restricted enhancer probes across 1,319 GSI-sensitive and 640 GSI-resistant allelic interactions showed a significant increase in their spatial perimeter (**Fig. S6i**, p = 0.002), suggesting expansion of the *LEF1* hyperinteracting hub in GSI resistance. This observation was in agreement with genomic data, which indicated the loss of an architectural stripe connecting *LEF1* with its flanking region (**Fig. S6j**).

Similar to the *LEF1* hub, optical mapping of the *IKZF2* locus using three-color Oligopaint DNA FISH with 3D confocal microscopy showed significant separation of the genomic elements at this hub in GSI-resistant DND41 (**Fig. 6f**, p = 0.003), in line with the visible loss of an architectural stripe on the *IKZF2* Hi-C contact frequency map (**Figure 6g**). Our single-cell resolution studies confirm the dynamic structure of enhancer-promoter hubs and further support their potential role in organizing regulation of genes involved in anticancer drug resistance.

### Ibrutinib resistance reorganizes enhancer-promoter hubs in MCL

The observation that enhancer-promoter hub spatial changes coincide with transcriptional changes associated with GSI resistance in T-ALL led us to examine whether the same relationship holds in MCL upon resistance to the BTK inhibitor ibrutinib, which is approved for the treatment of various hematological malignancies ^50^. We followed the same methodology as our T-ALL studies to identify differential hubs between ibrutinib-sensitive and ibrutinib-resistant Rec-1 MCL. Similar to GSI-resistant T-ALL, this analysis revealed that a small fraction of hubs (159 or ∼15%) were differential, notably gaining or losing interactivity in ibrutinib-resistant MCL (**Fig. 7a**). Similar to GSI-resistant T-ALL, differential hubs were separate from differential compartments and TADs identified in ibrutinib-resistant MCL (**Figs. S7a-d**), further supporting enhancer-promoter hubs’ status as distinct topological features (**Figs. S1a-c and S3a-c**). Investigation of epigenetic and transcriptional changes showed that significant loss or gain of interactivity in ibrutinib-resistant hubs coincided with marked decreases or increases in chromatin opening (**Fig. 7b**, p = 0.006; and **Fig. 7c**, p = 0.0009) and chromatin activity (**Fig. 7b**, p = 0.04; and **Fig. 7c**, p = 0.0004), respectively, with a similar trend for gene expression levels (F**ig. 7b**, p = 0.53; and **Fig. 7c**, p = 0.12).

**Figure 7:**
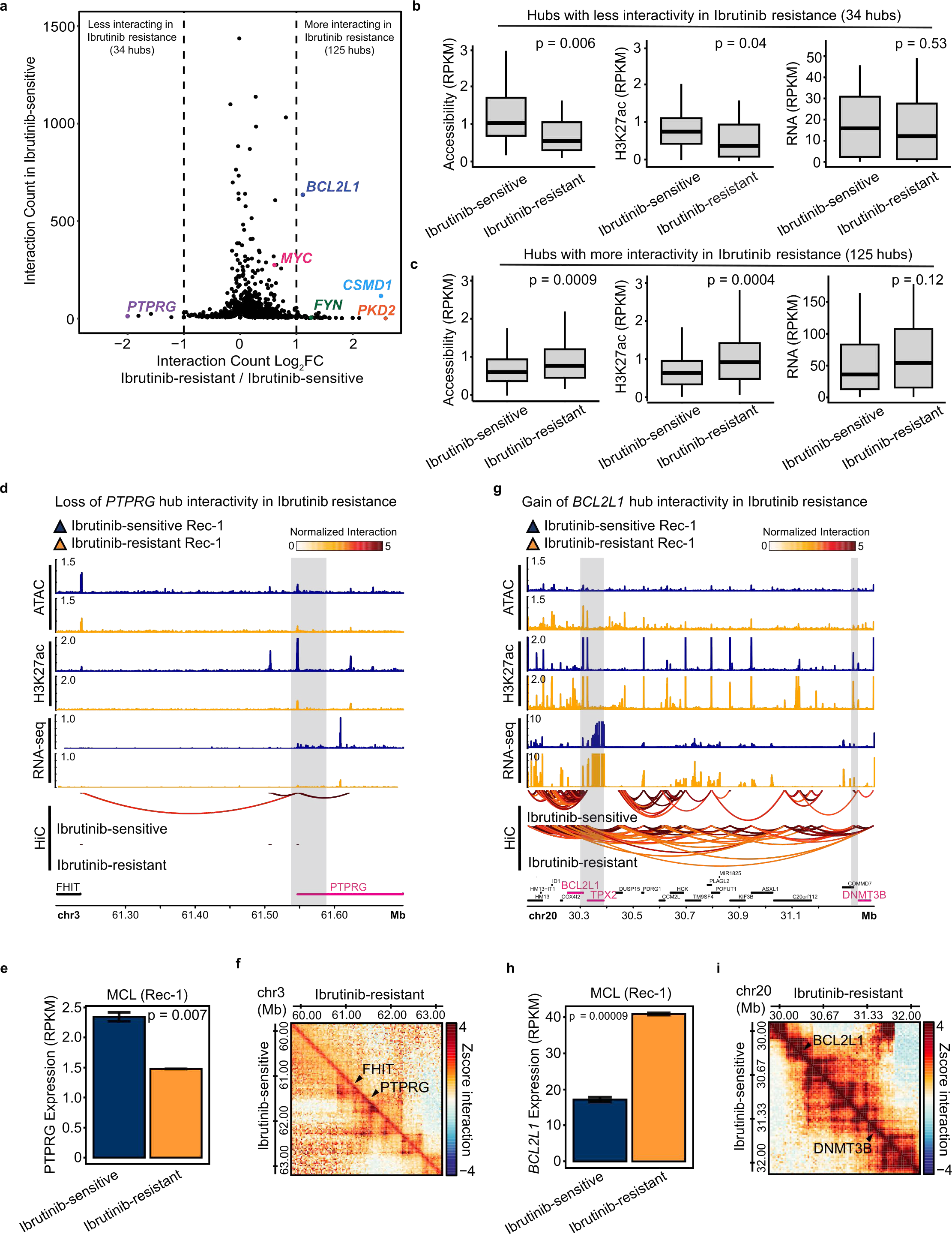
Genome-wide differential hub screen identifies loci that are aberrantly folded and expressed in ibrutinib-resistant MCL. a: Enhancer-promoter hubs with differential interactivity in ibrutinib-resistant vs. -sensitive MCL Rec-1 contain genes with potential roles in oncogenesis and / or treatment response. Scatter plot showing log_2_ fold change of hub interactivity in ibrutinib-resistant and -sensitive Rec-1 (x axis) versus hub interaction counts in ibrutinib-sensitive cells. Dotted lines mark enhancer-promoter hubs with more than or equal to 2-fold decrease (“less interacting in ibrutinib resistance”) or increase (“more interacting in ibrutinib resistance”) in interaction counts in ibrutinib-resistant vs. ibrutinib-sensitive Rec-1 cells. Hubs that were *de novo* gained or lost in ibrutinib resistance based on the hub minimum interactivity threshold of 6 were also considered differential. Selected hubs containing genes with potential role in oncogenesis and / or treatment response are labeled. For visual clarity, the two outlier hubs with >1,500 interaction counts in ibrutinib-sensitive and -resistant cells were excluded from the plot. b: Box-and-whisker plots showing significant reduction in chromatin accessibility (left) and H3K27ac deposition (center) as well as nonsignificant reduction in gene expression (right) of hubs with marked loss of interactivity in ibrutinib-resistant cells. Box-and-whisker plots: center line, median; box limits, upper (75th) and lower (25th) percentiles; whiskers, 1.53 interquartile range. P-value: two-tailed Wilcoxon rank sum test. c: Box-and-whisker plots showing significant increase in chromatin accessibility (left) and H3K27ac deposition (center), with a similar trend for transcription (right) in hubs that gained interactivity in ibrutinib-resistant cells. Box-and-whisker plots: center line, median; box limits, upper (75th) and lower (25th) percentiles; whiskers, 1.53 interquartile range. P-value: two-tailed Wilcoxon rank sum test. d: Genome tracks at the *PTPRG* locus show that loss of Hi-C interactions in the *PTPRG* hub in ibrutinib-resistant Rec-1 cells coincide with decrease of chromatin accessibility, H3K27ac active chromatin mark, and gene expression. Note that the arcs displayed in this illustration represent only the valid interactions detected from Hi-C between enhancer and/or promoter elements (with normalized contact frequency > 3). e: *PTPRG* is significantly downregulated in ibrutinib-resistant Rec-1 cells. Barplots of normalized RNA-seq reads showing *PTPRG* mRNA levels in ibrutinib-resistant and -sensitive cells. N: 3; P-value: unpaired student t-test; error bars: +/- 2 SE. f: Normalized Hi-C contact maps in ibrutinib-resistant (upper triangle) and ibrutinib-sensitive cells (lower triangle) at the *PTPRG* hub locus illustrates a marked loss in loops and subTADs connecting the *PTPRG* gene with surrounding elements in ibrutinib resistance. g: Genome tracks at the *BCL2L1* locus show that gain of Hi-C interactions in the *BCL2L1* hub in ibrutinib-resistant Rec-1 cells coincides with increase of chromatin accessibility, H3K27ac active chromatin mark, and transcription. Notice how two separate hubs in the ibrutinib-sensitive state combine to become a single hyperinteracting hub in the ibrutinib-resistant state. Note that the arcs displayed in this illustration represent only the valid interactions detected from Hi-C between enhancer and/or promoter elements (with normalized contact frequency > 3). h: *BCL2L1* is significantly upregulated in ibrutinib-resistant Rec-1 cells. Barplots of normalized RNA-seq reads showing *BCL2L1* mRNA levels in ibrutinib-resistant and -sensitive cells. N: 3; P-value: unpaired student t-test; error bars: +/- 2 SE. i: Normalized Hi-C contact maps in ibrutinib-resistant (upper triangle) and -sensitive cells (lower triangle) at the *BCL2L1* hub illustrates a marked gain in looping between *BCL2L1* and *DNMT3B (*as well as surrounding chromatin) in ibrutinib resistance.

Close inspection of hubs with loss of interactivity in ibrutinib-resistant MCL showed the dismantling of enhancer-promoter assemblies at genes involved in regulating GTPase activity, suggestive of dysregulation of GTPase signaling downstream of BTK inhibition within the BTK signaling cascade (**Table S2**) ^51, 52^. The hub containing tumor suppressor *PTPRG* ^53^ was the most differentially interacting hub with loss of interactivity in ibrutinib-resistant MCL (**Fig. 7d**). In line with loss of the *PTPRG* hub, we observed marked reductions in chromatin activity (**Fig. 7d**), Hi-C map contacts (**Fig. 7f**), and *PTPRG* expression levels at the *PTPRG* locus (**Fig. 7e**, p = 0.007).

In contrast, examination of hubs with gain of interactivity in ibrutinib-resistant MCL indicated that these topological assemblies predominantly formed at genes involved in processes promoting drug resistance, including cell cycle and apoptosis regulation, chromatin organization, stem cell maintenance, as well as regulation of protein transport (**Table S2**). Indeed, the hub containing *BCL2L1* (also known as *BCL-xL)*, a gene with proven anti-apoptotic function ^54^, gained over 700 interactions in ibrutinib resistance to become hyperinteracting (**Fig. 7g, Table S3**). Increases in chromatin activity over this hyperinteracting hub coincided with a significant increase in *BCL2L1* transcriptional activity (**Fig. 7h**, p = 0.00009) and visible changes in the Hi-C contact map (**Fig. 7i**), suggestive of a structural variation (such as a translocation) at this locus in ibrutinib-resistant MCL (**Table S4**). Other genes located in this hyperinteracting hub, including DNA damage response gene *TPX2* and DNA methyltransferase *DNMT3B*, were also markedly upregulated in ibrutinib resistance (**Fig. S7e**, p = 0.0004; and **Fig. S7f**, p = 0.0001). In line with observations in GSI-resistant T-ALL, this data further supports the role of enhancer-promoter hub reorganization in setting gene expression programs permissive to anticancer drug resistance.

## DISCUSSION

As an emerging unit of chromatin architecture, the spatial enhancer-promoter hub has remained a topological feature with cryptic structural and functional relevance. Here, we used a graph-based approach to systematically identify enhancer-promoter hubs from multi-omic data and examine their organizational principles and potential function in oncogenesis and drug resistance. By studying enhancer-promoter hubs in T-ALL, MCL, and TNBC, we found that these topological assemblies were distinct from TADs and ubiquitously enriched for transcription, with the most highly interacting hubs spatially coalescing genes encoding lineage-associated and oncogenic transcription factors and coregulators including *MYC*, *DOT1L*, *KAT5*, and *SOX9*, observations that may extend to other cancers.

Upon acquisition of Notch inhibitor resistance, a subset of hubs was reorganized as demonstrated by optical mapping of the *LEF1* and *IKZF2* hubs in individual T-ALL cells. Our data further showed that differential hubs contained genes that were recognized as key promoters of anticancer drug resistance in T-ALL ^21, 55^. Similarly, application of our differential hub screen to ibrutinib-sensitive and -resistant MCL revealed a variety of differential hubs containing genes with potentially important roles in supporting the drug-resistant phenotype, including *BCL2L1*. Indeed, it has been shown that *BCL2L1* inhibitors were able to induce potent cytotoxicity in MCL cells resistant to ibrutinib and *BCL2* inhibitor venetoclax ^56^. Although further investigation is required, our findings suggest that systematic examination of clusters of enhancers and promoters converging through space may guide identification of new therapeutic targets by revealing key regulatory elements and genes promoting drug resistance in cancer.

Our data also suggests that enhancer-promoter hubs uniquely straddle the structure-function axis. While the chain of causality remains elusive, our observations support a model of enhancer-promoter hubs in which the number of spatial interactions within a hub coincides with its relative transcriptional state. This model is reinforced by the correlation between transcriptional activity and enhancer positioning that has been documented at individual loci across various cancers ^14, 15, 57, 58^. It is likely that the mechanisms underlying enhancer-promoter hub formation also contribute to the distinct relationship between hub interactivity and transcription. Given the concordance of enhancer-promoter hubs identified from SMC1 HiChIP and Hi-C data, it is possible that cohesin-mediated loop extrusion may regulate hub spatial interactions and thus hub transcriptional activity, similar to cohesin’s role in organizing multi-way contact ‘hubs’ in single cells ^59^. In line with optical mapping performed in this study, further work is needed to concretely demonstrate the existence of multi-way interactions within hubs beyond correlative analysis of multi-omic data. Future studies leveraging advanced microscopy will refine our current understanding of hubs from population-based sequencing experiments and may shed light on hub landscapes within single cells. Taken together, our investigation suggests that enhancer-promoter hubs in cancer spatially organize transcriptional programs, which in turn may promote oncogenesis and drug resistance, parallel to enhancer-promoter hubs’ broader role in directing gene expression circuits in other diseases ^12, 13, 60, 61^.

## METHODS

### Contact For Reagent and Resource Sharing

Further information and request for reagents may be directed to and will be fulfilled by the corresponding author, Robert B. Faryabi.

### Experimental Procedures

#### Cell Culture

All of the data analyzed in this study for TNBC cell lines MB157 (ATCC, Cat# CRL-7721) and HCC1599 (ATCC, Cat# CRL-2331) and for T-ALL cell line CUTLL1 ^62^ were taken from previous investigations (see subsequent section). For the purpose of the FISH experiments conducted in this study, DND41 (DSMZ, Cat# ACC525) GSI-sensitive and GSI-resistant cells were cultured and maintained as previously described ^21^. Briefly, the DND41 cell line used for FISH analysis in this study was purchased from the Leibniz-Institute DSMZ-German Collection of Microorganisms and Cell Lines, and was grown in suspension with RPMI 1640 (Corning, cat# 10-040-CM) supplemented with 10% fetal bovine serum (Thermo Fisher Scientific, cat# SH30070.03), 2 mM L-glutamine (Corning, cat# 25-005-CI), 100 U/mL and 100 μg/mL penicillin/streptomycin (Corning, cat# 30-002-CI), 100 mM nonessential amino acids (GIBCO, cat# 11140-050), 1mM sodium pyruvate (GIBCO, cat# 11360-070) and 0.1mM of 2-mercaptoethanol (Sigma, cat# M6250). GSI-resistant DND41 cells were constantly cultured with GSI compound E (125 nM, Calbiochem, cat# 565790) and were periodically validated to have maintained the drug-resistant state by Western blotting for Notch intracellular domain 1 (NICD1), which is not present in cells constantly treated with GSI.

For experiments involving ibrutinib-sensitive and ibrutinib-resistant Rec-1 MCL, Rec-1 cells from the Genentech cell bank were cultured in RPMI 1640 (Corning, cat# 10-040-CM) supplemented with 10% fetal bovine serum (Thermo Fisher Scientific, cat# SH30070.03), 2 mM L-glutamine (Corning, cat# 25-005-CI), 100 U/mL and 100 μg/mL penicillin/streptomycin (Corning, cat# 30-002-CI), 100 mM nonessential amino acids (GIBCO, cat# 11140-050), 1mM sodium pyruvate (GIBCO, cat# 11360-070) and 0.1mM of 2-mercaptoethanol (Sigma, cat# M6250). Ibrutinib-resistant cells were generated over prolonged period of time using ibrutinib (Selleckchem, cat# S2680) dose escalation until they were stable in cell culture media supplemented with 100nM ibrutinib. Ibrutinib resistance was confirmed by 100-fold shift for ibrutinib IC50 between parental and resistant Rec1 lines. Parental and resistant cells, when used for RNA-seq, ATAC-seq, Hi-C, and ChIP-seq experiments, were treated with DMSO (parental) or ibrutinib (resistant) for 24 hours following BCR crosslinking with IgM. Cells used for ChIP-seq and Hi-C assays were subsequently fixed with 1% or 2% formaldehyde, respectively. Fixed frozen cell pellets were stored at -80C and used when needed.

#### Multi-omic Assays

In this study, we performed *in-situ* Hi-C, RNA-seq, H3K27ac ChIP-seq, and ATAC-seq on ibrutinib- sensitive and ibrutininb-resistant Rec-1 cells. Refer to the subsequent sections for descriptions on how these assays were performed. Excluding ibrutinib-sensitive and -resistant Rec-1 cells, multi-omic data for DND41, MB157, HCC1599, (untreated) Rec-1 cells, and CUTLL1 were obtained from previous investigations ^14, 21, 28, 29, 30^. Specifically, *in situ* Hi-C, SMC1 HiChIP, ATAC-seq, RNA-seq, H3K27ac ChIP-seq, and H3K27me3 CUT&RUN data from DND41 GSI-sensitive and GSI-resistant cells were downloaded from GSE173872. SMC1 HiChIP data for MB157, HCC1599, and Rec-1 were downloaded from GSE116876 along with H3K27ac ChIP-seq data for GSI-washout MB157, Rec-1, and HCC1599 and RNA-seq data for GSI-washout MB157 and HCC1599. RNA-seq data for GSI-washout Rec-1 was downloaded from GSE59810. GSI-washout cells were treated with GSI (1 uM, Calbiochem) for 72 hours before being washed and cultured in media containing DMSO for 5 hours, at which time RNA-seq and ChIP-seq assays were run. For the purposes of this study, data collected from GSI-washout cells

was considered equivalent to data collected from untreated cells for hub identification and analysis. For CUTLL1, *in situ* Hi-C and H3K27ac ChIP-seq data were downloaded from GSE115896, RNA-seq data was downloaded from GSE59810, and ATAC-seq data was downloaded from GSE216430.

#### Oligopaint FISH Probe Synthesis

DNA FISH Oligopaint probe libraries targeting three 50 Kb elements within the *LEF1* hub and three 75 Kb elements in the *IKZF2* hub were designed using OligoMiner ^63^. Each of the six distinct probe sublibraries was amplified and isolated from a larger pooled library using several short oligonucleotide primers (*RPL34* F primer: CTCGAATCGGTGTCGCATTC, R primer: TTGACGTTTGCGCCGAATAC; *LEF1* promoter F primer: TCCGCCGTGTTATCGATTTG, R primer: ATTCAACGGCCCTCGATTTG; *LEF1* enhancer F primer: TCATAATTCGGCGCTTGGTG, R primer: TGTATCGCGCGGTCAATTTC; *IKZF2* promoter F primer: TCGCTACGCCGGTTGTAATG, R primer: ATTACCGCGACCGGTTGAAG; *IKZF2* 5’ F primer: CAGTTACCGGTCCGTCGATG, R primer: ACGTATCGTCCCGCAACATG; *IKZF2* 3’ F primer: TTGTCGCGATGCCATAGACG, R primer: AGCTCAATCGTCGCACGATC), as previously described ^21^. Briefly, the pooled library was amplified via low cycle PCR and the T7 promoter was separately cloned into oligos within the six probe sublibraries of interest using primers specific to each probe sublibrary, which were purchased from IDT. Each probe was transcribed to RNA using a T7 RNA polymerase, and then RNA was reverse transcribed back into DNA oligos, which were purified and isolated for subsequent nuclear DNA hybridization. In order to visualize primary probes with fluorescence microscopy, secondary DNA probes conjugated with Alexa-488 (sequence: 5Alex488N/CACACGCTCTTCCGTTCTATGCGACG TCGGTG/3AlexF488N), Atto-565 (sequence: 5ATTO565N/ACACCCTTGCACGTCGTGGACCT CCTGCGCTA/3ATTO565N), and Alexa-647 (sequence: 5Alex647N/TGATCGACCACGGCCAA GACGGAGAGCGTGTG/3AlexF647N) reporters, which were also purchased from IDT, were used.

#### 3D Oligopaint DNA FISH on Slides

Cells were prepared for DNA FISH as previously described ^21^. Briefly, GSI-sensitive and GSI-resistant DND41 cells were first incubated on poly-*L*-lysine-coated glass slides (ThermoScientific, cat# P4981-001) and fixed in a solution of 4% formaldehyde in PBS for 10 minutes. Cell membranes were permeabilized by submerging slides in 0.5% Triton X-100 in PBS for 15 minutes and then denatured by immersion in increasing concentrations of ethanol (70%, 90%, and 100%). Cells were further permeabilized with cycles of immersion in heated 2x SSCT/50% formamide before incubation with a hybridization buffer containing 100 pmol of each of the three Oligopaint probes targeting the locus of interest. Slides were then sealed with rubber cement and a coverslip. After incubating in a 37C humidified chamber for 16 hours, the coverslip and probe hybridization solution were removed from the slides, which were once again cyclically submerged in 2x SCCT and 0.2x SCCT in heated water baths to re-permeabilize cell and nuclear membranes. Another hybridization mix containing secondary probes conjugated to fluorophores was aliquoted onto each slide before slides were sealed with rubber cement and coverslips. After incubation in the 37C humidified chamber for 2 hours, slides were submerged in 2x SCCT. DAPI dye was then added to stain nuclei, and slides were submerged in 2x SCCT for a final time. Finally, mounting media (Invitrogen, Ref# 336936) was added to each slide and coverslips were sealed onto the slides using transparent nail polish. Slides were then imaged using a Leica SP8 confocal microscope (*IKZF2* locus) with a 40x oil immersion objective or the Vutara VXL microscope (*LEF1* locus) (Bruker) in the widefield, epi-fluorescence microscope setting.

#### Rec-1 Ibrutinib *In-Situ* Hi-C

10^6^ parental or ibrutinib-resistant Rec-1 cells fixed with 2% formaldehyde were used as input for Hi-C assay performed with Arima Hi-C kit (Arima Genomics, cat#A510008) per the manufacturer’s instructions. Both samples passed Arima QC1 and were subsequently used for library generation with Accel-NGS 2S Plus DNA Library Kit (Swift, cat# 21024) and 2S Set A Indexing Kit (Swift, cat# 26148).

After passing Arima QC2, samples were PCR amplified for 6 cycles and quality was inspected using D5000 on Agilent 4200 Tapestation. Libraries were paired-end sequenced (2x150bp) using NovaSeq.

### Rec-1 Ibrutinib H3K27ac Chromatin Immunoprecipitation Sequencing (ChIP-seq)

ChIP-seq was performed using a previously published protocol ^14^. 10x10^6^ parental or ibrutinib-resistant Rec-1 cells previously fixed with 1% formaldehyde (Pierce, cat#28908) and quenched with 0.125M Glycine (Fisher scientific, cat#AAJ1640736), were sonicated for 5.5 min on Covaris L220 with the following settings: PIP 350, DF 15%, CPB 200. Chromatin was cleared with recombinant protein G– conjugated agarose beads (Invitrogen, cat# 15920-010) and subsequently immunoprecipitated with H3K27ac antibody (Active Motif, cat# 39133). Antibody-chromatin complexes were captured with recombinant protein G–conjugated agarose beads, washed with Low Salt Wash Buffer, High Salt Wash Buffer, LiCl Wash Buffer and TE buffer with 50mM NaCl and eluted. Input sample was prepared by the same approach without immunoprecipitation. After reversal of cross-linking, RNase (Roche, cat# 10109169001) and Proteinase K (Invitrogen, cat# 25530-049) treatments were performed, and DNA was purified with QIAquick PCR Purification Kit (QIAGEN, cat# 28106). Libraries were then prepared using the NEBNext Ultra II DNA library Prep Kit for Illumina (NEB, cat# E7645S). Two replicates were performed for each condition. Indexed libraries were validated for quality and size distribution using a TapeStation 4200 (Agilent). Libraries were paired-end sequenced (2x50bp) on Illumina NextSeq instrument.

#### Rec-1 Ibrutinib Assay for Transposase Accessible Chromatin (ATAC-seq)

ATAC-seq assay was performed as previously described ^14^. Briefly, 60,000 parental or ibrutinib-resistant Rec-1 cells were washed with 50 μL of ice cold 1 x PBS (Corning, cat# 21031CV), followed by 2 minute treatment with 50 μL lysis buffer containing 10 mM Tris-HCl, pH 7.4, 3 mM MgCl_2_, 10 mM NaCl, and 0.1% NP-40 (Igepal CA-630). After pelleting nuclei, nuclei were resuspended in 50 μL of transposition buffer consisting of 25 μL of 2x TD buffer, 22.5 μL of molecular biology grade water, and 2.5 μL Tn5 transposase (Illumina, cat# FC-121-1030) to tag the accessible chromatin for 45 minutes at 37C. Tagmented DNA was purified with MinElute Reaction Cleanup Kit (QIAGEN, cat# 28204) and amplified with 5 cycles. Additional number of PCR cycles was determined from the side reaction and ranged from 8-9 total cycles of PCR. Two replicates were performed for each condition. Libraries were purified using QiaQuick PCR Purification Kit (QIAGEN, cat# 28106) and eluted in 20 μL EB buffer. Indexed libraries were assessed for nucleosome patterning on the TapeStation 4200 (Agilent) and paired-end sequenced (50bp+50bp) on HiSeq (Illumina).

#### Rec-1 Ibrutinib RNA Sequencing (RNA-seq)

Strand-specific RNA-seq was performed on parental and ibrutinib-resistant Rec-1 cells using SMARTer Stranded Total RNA Sample Prep Kit (Takara, cat# 634873) per the manufacturer’s instructions. Briefly, 24 hours post treatment with DMSO and 100nM ibrutinib followed by IgM stimulation for 5 min, parental and resistant cells were lysed with 350 μL RLT Plus buffer (QIAGEN) supplemented with 2-mercaptoethanol (Sigma, cat# M6250) and total RNA was isolated using the RNeasy Plus Micro Kit (QIAGEN, cat# 74034). RNA integrity numbers were determined using TapeStation 2200 (Agilent). 800 ng of total RNA was used, and libraries were prepared using the SMARTer Standard Total RNA Sample Prep Kit - HI Mammalian. Libraries were paired-end sequenced (38+38 bp) on a HiSeq. Three biological replicates were performed for each condition.

### Quantification and Statistical Analysis

#### Definition of Regulatory Elements

For the purposes of this study, regions within 2.5 Kb of transcription start sites (TSSes) of expressed genes were considered to be promoters. H3K27ac ChIP-seq peaks that did not overlap with promoter

regions were considered to be enhancers. For identification of hubs from Hi-C data, only enhancers and promoters that overlapped with ATAC-seq peaks were considered valid loop anchors.

#### Gene Annotation

All genes presented in this study were annotated in accordance with the Human Genome version 19 (hg19) GRCh37.75 assembly. 2,828,317 Ensembl transcripts from this assembly were downloaded, and the longest transcript from each Ensembl gene id (ENSG) was used to generate a list of transcription start sites (for promoter annotation) and gene position. 57,209 gene annotations were used for RNA-seq analysis after excluding rRNA and chrM genes.

#### Rec-1 Ibrutinib H3K27ac ChIP-seq and ATAC-seq Data Analysis

Rec-1 ibrutinib-sensitive and -resistant H3K27ac ChIP-seq reads were trimmed with Trim Galore (version 0.6.6) using parameters -q 5 --phred33 --fastqc --gzip --stringency 5 -e 0.1 --length 20 –paired. Ensembl GRCh37.75 primary assembly, which included chromosome 1-22, chrX, chrY, chrM and contigs was used for alignment of trimmed reads with BWA (version 0.7.13) ^64^. BWA was run using the command bwa aln -q 5 -l 32 -k 2 -t 12 and paired-end reads were grouped using bwa sampe -P -o 1000000 -r. After grouping, reads which were considered duplicates from Picard (version 2.1.0) as well as reads that were matched with ENCODE blacklist regions or contigs were filtered out so that valid reads were kept and used for all subsequent analysis. This procedure was repeated for alignment of Rec-1 ibrutinib-sensitive and -resistant ATAC-seq reads. See the following section for ChIP-seq and ATAC-seq peak-calling protocols.

#### Rec-1 Ibrutinib Hi-C Data Analysis

HiC-Pro (version v2.8.1) ^65^ was used to process Rec-1 ibrutinib-sensitive and -resistant Hi-C raw reads for each sample using default parameters except LIGATION_SITE and GENOME_FRAGMENT were provided by Arima. Putative interactions identified from HiC-Pro were used as the basis for hub calling. Similar to Rec-1, CUTLL1 Hi-C data was processed using HiC-Pro (version v2.8.1) with default parameters.

#### H3K27ac ChIP-seq and ATAC-seq Peak Calling

Peak calling of all H3K27ac ChIP-seq and ATAC-seq data was performed similar to as previously described ^14^. Briefly, fragment length of H3K27ac ChIP-seq reads was estimated with HOMER (version 4.8) ^66^, and MACS was used to identify peaks with the parameters -q 1E-5 –shiftsize = 0.5∗fragment_length –format = BAM –bw = 300 –keep-dup = 1. After peak calling, H3K27ac signal over peaks was quantified and normalized to reads per kilobase per million mapped reads (RPKM). ATAC-seq reads were processed similarly to H3K27ac reads except MACS peak calling was performed with parameters -p 1E-5 –nomodel –nolambda –format = BAM –bw = 300 –keep-dup = 1. After peaks were called for each condition, BED files containing the union of peaks across relevant conditions (e.g. drug-sensitive, drug-treated, and drug-resistant) were created using the bedtools merge command. Note that for CUTLL1 H3K27ac ChIP-seq and ATAC-seq data, a slightly less stringent q value cutoff of 1E-4 was used for peak calling to obtain a comparative number of peaks between cell lines. Exact steps for each workflow can be found at https://github.com/faryabiLab/dockerize-workflows/tree/master/workflows.

#### RNA-seq Analysis

Bulk total transcript RNA-seq data from T-ALL DND41 and CUTLL1, MCL Rec-1 (including ibrutinib-sensitive and -resistant), and TNBC MB157 and HCC1599 cells was used to annotate expressed genes and analyze RNA enrichment over hub intervals. Alignment to Ensembl GRCh37.75 primary assembly and normalization with RPKM was performed as previously described ^21^. RNA-seq for DND41, Rec-1, MB157, and HCC1599 was originally performed on at least three replicates for each cell line, and expressed genes were determined using a cutoff of RPKM > 1 in at least two of three replicates. For CUTLL1, a single RNA-seq experiment was analyzed with expressed genes determined using the same cutoff of RPKM > 1.

#### Topologically Associating Domain (TAD) Boundary and Differential TAD Boundary Identification

TAD boundaries were identified in both DND41 and Rec-1 drug-sensitive and drug-resistant cells from SMC1 HiChIP and Hi-C data as previously described ^21^. Briefly, the cooltools insulation function (https://cooltools.readthedocs.io/en/latest/) was applied to a .cool file converted from HiC-Pro ^65^ valid pairs output using the hicpro2higlass.sh function followed by the hic2cool command-line tool with default options. Using a window size of 100 Kb and a bin size of 5 Kb, insulation scores were calculated. Valid, adjacent boundaries were defined as those with total reads between them exceeding the 75th percentile. Differential Hi-C TAD boundaries between drug-sensitive and drug-resistant conditions were categorized as boundaries with absolute log2 fold change in insulation score greater than 0.75.

#### Differential Compartment Identification

Hi-C data was used to detect differential compartments between drug-sensitive and drug-resistant DND41 and Rec-1. Compartments were initially identified in each condition from the first principle component (PC1) of Hi-C data, which was calculated with the Homer v4.11 ^66^ runHiCpca.pl method using H3K27ac ChIP-seq data to avoid arbitrary sign assignment. Next, the getHiCcorrDiff command was used to calculate each compartment’s correlation difference between drug-sensitive and drug-resistant conditions. To identify differential compartments switching from A to B in the drug-resistant state, the findHiCCompartments command was used with a correlation difference threshold (-corr) and a PC1 threshold of at least 0.4 for DND41 and 0.65 for Rec-1. For detection of compartments switching from B to A in the drug-resistant state, the same findHiCCompartments command was used with the addition of the -opp flag.

### Enhancer-Promoter Hub Identification and Analysis

#### Spatial interaction pre-processing overview

Enhancer-promoter hubs in T-ALL DND41, T-ALL CUTLL1, MCL Rec-1, TNBC MB157, and TNBC HCC1599 were identified using Hi-C or SMC1 HiChIP contact frequency data annotated for enhancers and promoters. Specifically, SMC1 HiChIP data was used for detection of hubs in DND41, HCC1599, MB157, and Rec-1 cells whereas Hi-C data was used for detection of enhancer-promoter hubs in CUTLL1, GSI-sensitive and -resistant DND41 cells and ibrutinib-sensitive and -resistant Rec-1 cells. A high level description of the data filtering process to create the inputs for our hub identification program is as follows: in order to generate data tables of valid enhancer-enhancer (EE), enhancer-promoter (EP), or promoter-promoter (PP) interactions in each condition, Hi-C/HiChIP loop anchors were first filtered by intersecting them with BED files of H3K27ac ChIP-seq peaks (i.e. enhancers) and active transcription start sites (TSS) from RNA-seq (i.e. promoters). For this analysis, peaks were defined as the 5 Kb or 10 Kb centered around the summit of the original protein enrichment signal. Anchors that were annotated as active TSSes with H3K27ac peaks were considered to be promoters. Once spatial interactions between putative promoter and/or enhancer elements were isolated, they were assigned normalized contact frequency scores and further filtered (using a user-defined score cutoff for Hi-C) to create the final data table for input into the hub-calling algorithm. This process of interaction filtering was slightly different depending upon the source assay of contact frequency data (i.e. Hi-C vs. SMC1 HiChIP) and upon other genomic data (e.g. ATAC-seq) that was readily available for analysis.

#### SMC1 HiChIP spatial interaction pre-processing

For SMC1 HiChIP data, valid EE/EP/PP input interactions into the hub-calling pipeline were detected from raw reads by filtering significant interactions identified from FitHiChIP v11.0 ^67^ to only include those between enhancers and active promoters. Briefly, SMC1 HiChIP reads were processed with Hi-C

Pro version v2.5.0 ^65^. High-confidence loop anchors were identified from FitHiChIP using Hi-C Pro’s allValidPairs file as input, a significance cutoff of q = 0.05 or p = 0.05 (see next paragraph), coverage bias regression for normalization, an interaction type of all to all / IntType=4, and default values for the remaining options. Anchors of significant FitHiChIP interactions were divided into two BED files and separately intersected with a BED file containing the union of H3K27ac ChIP-seq peaks and actively transcribed TSSes, where active genes were defined as those with >1 RNA-seq RPKM in at least two thirds of the replicates across the conditions of interest. After valid promoter and enhancer anchors were identified, the original list of FitHiChIP interactions was filtered to keep only interactions between valid enhancer and/or promoter elements for input to the hub-calling pipeline.

For filtering interactions from Rec-1, MB157, and HCC1599 with FitHiChIP, a significance threshold of q < 0.05 was used to yield over 100,000 significant interactions in each cell type. However, since FitHiChIP identified less than 25,000 significant interactions from DND41 SMC1 HiChIP data with a significance cutoff of q < 0.05, we decided that a significance threshold of p < 0.05 was more appropriate for filtering interactions in this cell line. To ensure stringent filtering of DND41 SMC1 HiChIP data given this lower threshold, FitHiChIP significant interactions (p < 0.05) for DND41 were subject to further filtering. Specifically, BED files containing anchors of significant FitHiChIP interactions, ATAC-seq peaks, SMC1 ChIP-seq peaks, and the union of H3K27ac ChIP-seq peaks and actively transcribed TSSes were intersected to create a list of high confidence, accessible regulatory element anchors. Only significant FitHiChIP interactions between two of these anchors were considered valid and used for DND41 hub calling. Since enhancers were not filtered by accessibility in Rec-1, MB157, and HCC1599 cells, they were defined more stringently as H3K27ac ChIP-seq peaks of > 500 bp.

#### In-situ Hi-C spatial interaction pre-processing

For Hi-C data, valid EE/EP/PP interactions were filtered from Hi-C reads by quantifying the number of contacts between accessible enhancers and accessible, actively expressed promoters. First, a data table of all possible *cis* interactions between accessible enhancers and accessible, active promoters was generated for the conditions of interest. BED files containing active, accessible promoters were created by intersecting a BED file of actively transcribed TSSes, where active genes were defined as those with >1 RNA-seq RPKM in 2 out of 3 replicates for either drug-sensitive *or* drug-resistant cells, with a BED file containing the union of ATAC-seq peaks from drug-sensitive and drug-resistant conditions. BED files of accessible enhancers were created by intersecting a BED file containing the union of H3K27ac ChIP-seq peaks from drug-sensitive and drug-resistant cells with a BED file containing the union of ATAC-seq peaks from drug-sensitive and drug-resistant cells. A matrix containing all possible combinations of *cis* linkages (maximum length of 2 Mb) between these accessible enhancer and promoter elements was then constructed and used to normalize Hi-C reads for a given condition. Specifically, Hi-C reads were processed with Hi-C Pro ^65^, rearranged into BEDPE file format, and intersected with the aforementioned BED file of ATAC-seq union peaks such that only interactions between pairs of accessible anchors were kept. These contacts were then mapped onto the matrix containing all possible enhancer and promoter interactions, and the number of ATAC-filtered Hi-C reads overlapping with each possible EE/EP/PP interaction was counted. Next, these linkage counts were summed and normalized to contacts per 100 million, and interactions with a normalized contact frequency of >3 (for DND41 and Rec-1) or >5 (for CUTLL1) were considered valid interactions for hub calling.

#### Hub-calling pipeline

We adapted an implementation of matrix-free divisive hierarchical spectral clustering (https://github.com/faryabiLab/hierarchical-spectral-clustering), which was originally developed for single cell RNA-seq ^20^, to identify enhancer-promoter hubs from the enhancer-promoter connectivity graph. We reasoned that this approach can overcome limitations associated with heuristic global optimization-based community detection methods such as Louvain-based algorithms to efficiently identify enhancer-promoter hubs, which we define as groups of densely connected enhancers and promoters with high intra-group and sparse inter-group interactions. Briefly, our matrix-free divisive hierarchical spectral clustering uses all of the information embedded in the enhancer-promoter connectivity graph at each partitioning to create a tree of enhancer and/or promoter clusters by recursively bi-partitioning the input spatial interactions between genomic elements. Time and memory efficiency is achieved by replacing factorization of the normalized Laplacian matrix at each iteration with direct calculation of the second left singular vector corresponding to the second largest singular value of a new matrix derived from the sparse connectivity matrix ^20^. To enable simultaneous detection of large and small interacting enhancer/promoter clusters and to avoid creating arbitrary small clusters, our approach uses Newman-Girvan modularity ^68^ as a stopping criterion for recursive cluster bi-partitioning. Using modularity as a stopping criterion instead of an optimization parameter also bypasses limitations associated with heuristic global optimization-based clustering such as Louvain-based algorithms ^69, 70^.To this end, our approach produces a hierarchy of nested enhancer-promoter clusters where each inner node is a cluster at a given scale and each leaf node is the finest-grain cluster such that any additional partitioning would be as good as randomly splitting connected enhancers/promoters. Importantly, this divisive hierarchical spectral clustering approach maintains relationships among clusters at various levels. As a result, the nested cluster structure can be used for clustering tree pruning, which provides flexibility in analysis and interpretation of enhancer-promoter hubs when hub topologies are *a priori* unclear.

To prevent the possibility that a functionally relevant cluster is divided into multiple clusters for downstream analysis, we used parameters cluster-tree -c dense -n 2 -s for all the analysis, and defined hubs as self-contained networks of connected regulatory elements. We next categorized hubs by the largest contiguous genomic interval covered by their enhancer/promoter anchors, and computed their within-hub spatial interaction counts and enhancer/promoter counts. We also annotated hubs with the expressed genes contained within their contiguous intervals (**Table S1**). To ensure that we identified hubs rather than just a few interacting elements, we removed clusters containing fewer than 6 spatial interactions among less than 4 enhancer/promoters for downstream analysis. For each cell type, distributions of hubs were plotted on the basis of their interaction count and enhancer/promoter count. The interaction count cutoff for hyperinteracting hubs was determined by calculating the point of tangency on the elbow of the curve comparing hub rank to interaction count as previously described ^14^.

#### Hub vs. random loci RNA enrichment analysis

Scatterplots of median RNA enrichment over hub loci and random representative hub loci were created for each cancer type to analyze transcriptional activity of hubs. Lists of random representative hub loci were generated by iteratively calling the bedtools shuffle command with parameters -chrom - noOverlapping -g hg19.genome on input BED files containing observed hubs. This bedtools command generates a list of random loci that contains an identical number of loci as the input list of observed hubs, where each random locus on the output list has identical length and chromosomal localization as each observed hub in the input list. As noted in the bedtools documentation, the process of generating random representative loci using this command could very infrequently skip loci violating the noOverlapping criteria. Median RNA enrichment was then quantified over each list of random representative loci to create a single data point on the output scatterplot, and this process was repeated several thousand times (5,000 times for all total hub analyses and 10,000 times for all hyperinteracting hub analyses) to enhance the accuracy of the simulated comparison.

#### SMC1 HiChIP vs. Hi-C hub similarity Venn diagrams

In order to compare DND41 and Rec-1 hubs identified from Hi-C with DND41 and Rec-1 hubs identified from SMC1 HiChIP (respectively), Venn diagrams were used to depict the similarity of hubs on the basis of their genomic overlap. These diagrams indicate the percentage of distinct, non-overlapping HiChIP hubs (relative to the total number of HiChIP hubs) and Hi-C hubs (relative to the total number of Hi-C hubs) as well as the percentages of overlapping HiChIP and Hi-C hubs reported both in terms of the total number of HiChIP hubs and the total number of Hi-C hubs. Note that the sizes of the Venn diagram circles are proportional to the number of hubs between those conditions.

#### All hub and hyperinteracting hub similarity matrices

Similarity matrices of all hubs and hyperinteracting hubs identified from Rec-1, HCC1599, MB157, and DND41 SMC1 HiChIP were created by comparing the genomic positions of hubs between each pair of cancer types. For example, to compare SMC1 HiChIP hyperinteracting hubs between DND41 and Rec-1 cells, the number of non-overlapping (where overlap is defined as ≥ 1bp) hyperinteracting hubs in each cancer was calculated by using bedtools intersect –v on two input BED files of hyperinteracting hubs in each cancer type. The similarity score for the comparison was then calculated using the following formula: similarity = 100 – (average percentage of distinct hubs). Since no more than 5% of within-condition hubs or hyperinteracting hubs for SMC1 HiChIP were overlapping, this methodology of similarity analysis was considered to be minimally biased.

The ten shared (i.e. overlapping) hyperinteracting hubs across T-ALL DND41, MCL Rec-1, and TNBC MB157 and HCC1599 were identified by finding the intersection of the four SMC1 HiChIP hyperinteracting hub lists using bedtools intersect, and manually filtering the resulting overlapping intervals based upon their connectivity structure and size. Specifically, overlapping regions were only considered to be conserved hyperinteracting hubs if there were valid SMC1 HiChIP contacts over the overlapping region in all four cell lines and if the region spanned more than ∼75 Kb (i.e. size of the smallest hyperinteracting hub across the four cell types).

#### TAD and hub boundary similarity analysis

Hub boundaries were defined as the 5 Kb upstream and downstream from the start position and end position of hubs (respectively) resulting in 2h intervals, where h is the number of hubs identified. Valid TAD boundaries from DND41 and Rec-1 SMC1 HiChIP or Hi-C data were identified as described in the previous section and intersected with the aforementioned HiChIP or Hi-C hub boundaries (respectively) to produce Venn diagrams depicting hub/TAD boundary overlap. Additionally, pile-up plots of Hi-C hub and Hi-C TAD boundaries were separately generated using coolpup.py with options –pad 250000 – local and graphed with plotpup.py ^71^. Note that few boundaries in each condition were automatically excluded from pile-up plots by coolpup.py.

#### Hyperinteracting vs. regular hub overlap with super-enhancers

Super-enhancers were identified in each cell line by applying previously described methods ^72^ to H3K27ac ChIP-seq peaks for CUTLL1, DND41, Rec-1, MB157, and HCC1599 in R. Super-enhancers were intersected with hub genomic locations and stratified by type (hyperinteracting or regular). Super-enhancer and hub similarity was determined by calculating the proportion of regular and hyperinteracting hubs overlapping with 0, 1, or 2+ super-enhancers.

#### Hyperinteracting vs. regular hub genomic length violin plots

Genomic lengths of hubs were calculated as the difference between the start coordinate of the farthest upstream regulatory element in the hub and the stop coordinate of the farthest downstream regulatory element in the hub for DND41, Rec-1, MB157, and HCC1599 hubs identified from SMC1 HiChIP data. These genomic distances were stratified by hub type (i.e. hyperinteracting vs. regular) and plotted in R using ggplot2’s geom_violin function.

#### Hyperinteracting vs. regular hub highly expressed gene plots

Highly expressed genes were considered to be genes within the top 2.5% quantile of gene expression, as defined from RNA RPKM in DND41, Rec-1, MB157, and HCC1599. The genomic coordinates of these highly expressed genes were intersected with hyperinteracting and regular hubs, and the number of highly expressed genes overlapping each hyperinteracting or regular hub was plotted in R using ggplot2’s geom_violin function.

#### Identification of structural variants in MCL

Structural variations in ibrutinib-sensitive and ibrutinib-resistant Rec-1 were identified from ICE-balanced Hi-C data using the predicts command of EagleC v0.1.9 with default options ^73^. Hi-C matrices were prepared as .cool files with resolutions 10 Kb, 50 Kb, and 100 Kb, as required by the EagleC pipeline.

### Differential Enhancer-Promoter Hub Identification and Analysis

#### Differential hub-calling pipeline

Differential enhancer-promoter hubs were identified from the hub-calling output for two different conditions (i.e. two BED-like files containing hub genomic intervals and interaction counts). In order to compare hubs in each condition on the basis of interaction count, the bedtools merge command was used to find the union of hubs across the two conditions. For union hubs that were created from ≥ 3 overlapping precursor hubs, new interaction counts were calculated by summing the number of interactions of the individual precursor hubs in each condition. The union hub lists also contained *de novo* lost/gained non-overlapping hubs. Finally, the log_2_ fold change in interaction count over union hubs was calculated, where log_2_ fold change = log_2_(case interaction count) - log_2_(control interaction count), and union hubs were annotated by the expressed genes contained within their contiguous genomic intervals.

#### Differential hub analysis

For each set of union hubs, a volcano-like plot comparing the drug-sensitive interaction counts and log_2_ fold change in interaction count of each union hub was graphed. These plots were used to illustrate genome-wide changes in hub interactivity, and to highlight large differential hubs of interest for downstream analysis. Hubs were considered to become more interacting in the drug-resistant state either if log_2_FC(interaction count) ≥ 1 or if they increased from < 6 interactions to ≥ 6 in the drug-resistant condition (i.e. *de novo* gained hubs). Similarly, hubs were considered to become less interacting in the drug-resistant state either if log_2_FC(interaction count) ≤ -1 or if they decreased from ≥ 6 interactions to < 6 interactions in the drug-resistant condition (i.e. lost hubs). In order to best visualize all of the hubs in the volcano-like plot for DND41 GSI-sensitive vs. GSI-resistant hubs, the log_2_ fold change of all hubs in the plot was adjusted using a pseudocount such that log_2_ fold change = log_2_[(GSI-resistant interaction count + 1) / (GSI-sensitive interaction count + 1)]. This pseudocount adjustment was not necessary to create the volcano-like plot of differential hubs in ibrutinib-sensitive vs. ibrutinib-resistant Rec-1. RNA enrichment, H3K27ac enrichment, and ATAC-seq enrichment were then examined over these differential hubs. Gene ontology and pathway enrichment analysis over the expressed genes contained within differential hubs was also performed.

#### Differential compartment and differential hub similarity Venn diagrams

Differential A to B and B to A compartments were identified as described in the previous section and intersected with differential hubs that lost or gained interactivity in the drug-resistant condition, respectfully. For similarity analysis, the overlap between the full genomic length of differential hubs and differential compartments was calculated and plotted with Venn diagrams.

#### Differential TAD and differential hub boundary similarity Venn diagrams

Differential TAD boundaries losing insulation or gaining insulation were identified as described in the previous section and intersected with boundaries of differential hubs that lost or gained interactivity in the drug-resistant condition, respectfully. For similarity analysis, the overlap between the differential hub and differential TAD boundaries was calculated and plotted with Venn diagrams.

### Genomic Feature Intersection/Similarity Analyses

Intersection analysis to determine the linear genomic overlap between distinct populations of enhancer-promoter hubs and between hubs and other genomic features (e.g. TADs, compartments, super-enhancers, etc.) was performed using either the intersect command of bedtools ^74^ or the findOverlaps command of R library GenomicRanges. Unless otherwise noted, for hub populations and/or genomic features that overlapped such that one hub in list A intersected more than one hub/feature in list B (or vice versa), bedtools was used to determine the degree of overlap between the two lists. For genomic features and hubs that overlapped such that at most one hub in list A intersected one feature in list B (and vice versa), GenomicRanges was used to determine the degree of overlap between the two lists.

All Venn diagrams illustrating the output of feature intersection analysis were created in R using the eulerr package such that each segment’s size was proportional to the number of features that it represented. For Venn diagrams illustrating intersection analyses that had at least one feature in one list intersecting more than one feature in the other, the sizes of the diagram segments were set such that they reflected the number of exclusive features in each set and the average degree of overlap, where average overlap = (total number features - total number exclusive features)*0.5. All Venn diagram segments were manually labeled with the actual number and percentage of exclusive and overlapping features in each list.

### Gene Ontology (GO) and Pathway Enrichment Analysis

GO enrichment analysis was conducted using the enrichment function of Metascape ^75^ or the PANTHER enrichment function of the Gene Ontology Knowledgebase ^76, 77, 78^. For hyperinteracting hubs in each condition, the list of expressed genes contained within the intervals of hyperinteracting hubs was used as input for analysis of GO molecular function, biological process, and/or cellular component annotation enrichment. Metascape was used for GO enrichment analysis of SMC1 HiChIP hyperinteracting hubs in Rec-1, HCC1599, and MB157 while PANTHER was used for GO enrichment analysis of SMC1 HiChIP hyperinteracting hubs in DND41 and Hi-C hyperinteracting hubs in CUTLL1 because the number of expressed genes in T-ALL hyperinteracting hubs exceeded the gene limit for Metascape. Metascape was also used for GO enrichment analysis of DND41 and Rec-1 hyperinteracting hubs identified from Hi-C data. For GO enrichment analysis of differential DND41 and Rec-1 hubs, expressed genes in drug-resistant cells that were located within hubs that gained interactivity or expressed genes in drug-sensitive cells that were located within hubs that lost interactivity were used as inputs to Metascape. In addition to GO analysis, pathway enrichment analysis was also performed on these expressed genes within differential hubs in Metascape using the Hallmark Gene Set, Reactome Gene Set, Biocarta Gene Set, Canonical Pathway, WikiPathway, and KEGG Pathway annotations.

For discussion of Metascape GO/pathway enrichment output in this paper, significant ‘summary’ annotations (p < 0.01) were presented (as opposed to significant ‘non-summary’ annotations) given that the ‘summary’ annotations represented overarching groups of GO terms and therefore limited redundancy in annotation reporting. For all Metascape hyperinteracting hub GO plots (Figs. 3g, 4g, and S4m), the top 10 most significantly enriched ‘summary’ GO molecular function annotations from expressed genes in SMC1 HiChIP hyperinteracting hubs were ranked by p-value and plotted. For presentation of PANTHER GO enrichment output, we plotted the top 10 most significantly enriched GO molecular function annotations ranked by p value, excluding selected annotations from the plots. More specifically, Figs. 2g and S2h show top 10 GO terms excluding the highly enriched ‘unclassified’ and ‘molecular function’ GO terms because these are not informative. Also note that the ‘binding’ GO term in these figures is not immediately relevant for direct gene annotation given its breadth as a parent GO annotation.

### DNA FISH Analysis

DNA FISH image analysis was performed similar to a previously described protocol ^21^. Briefly, DAPI signal was used for manual nuclei segmentation, with 1548 GSI-sensitive allelic interactions and 533 GSI-resistant allelic interactions analyzed for the *IKZF2* locus and 1319 GSI-sensitive allelic interactions and 640 GSI-resistant allelic interactions analyzed for the *LEF1* locus. For each manually segmented nucleus, ‘spots’ indicative of probe signal were manually thresholded. Centroid positions for each spot in xy were found by fitting a Gaussian. X, Y, and Z coordinates were extracted, and pairwise Euclidean distances between nearest neighbors were calculated. Representative FISH cell images were taken using a Leica SP8 63x oil immersion objective and photo brightness, contrast, and smoothing was adjusted in ImageJ to facilitate probe visualization.

### Genomic Data Visualization

Normalized reads from ATAC-seq, H3K27ac ChIP-seq, H3K27me3 CUT&RUN, RNA-seq, SMC1 HiChIP, and/or Hi-C over selected hubs were visualized using the R package Sushi (version 1.18.0) ^79^. Specifically, bedgraph (.bg) files of reads normalized using reads per million from ATAC-seq, H3K27ac ChIP-seq, H3K27me3 CUT&RUN, and RNA-seq data were created with command genomecov, or by converting RPM-normalized BigWig (.bw) files into .bg format using the UCSC tools (version 329) BigWigToBedGraph ^80^. These .bg files were visualized using the Sushi command plotBedgraph. For Hi-C data, normalized EE/EP/PP interactions exceeding the contact frequency cutoff of 3 (for DND41 and Rec-1) or 5 (for CUTLL1) were plotted using the Sushi command plotBedpe with intensity equivalent to normalized contact frequency. For SMC1 HiChIP data, EE/EP/PP FitHiChIP significant interactions (p < 0.05 for DND41 and q < 0.05 for Rec-1, MB157, and HCC1599) were plotted with intensity equivalent to –log_10_(X), where X is the minimum FitHiChIP p-value (for DND41) or q-value (for Rec-1, MB157, and HCC1599) for the interaction. Z-scored contact maps were created to visualize Hi-C interaction heatmaps over selected hubs. These plots were generated by first applying z-score transformation to 25Kb resolution VC_SQRT normalized Hi-C contact maps for each chromosome as previously described ^81^ using the R command loessFit with parameters iter = 100, span = 0.02. The Sushi command plotHiC was used to plot these transformed maps.

### Data Presentation & Statistical Analysis

All analysis and quantification of ATAC-seq and ChIP-seq output was performed using peak-called data with normalized RPKM measurements. Median RNA enrichment over observed spatial hubs and hyperinteracting hubs was compared to median RNA enrichment over random representative loci using empirical p-values conservatively calculated as (n+1)/(r+1), where r is the total number of replicates of random representative loci lists (i.e. 5,000 or 10,000) and n is the number of replicates in which theoretical median RNA enrichment exceeded observed median RNA enrichment over hubs/hyperinteracting hubs. Two-tailed Wilcoxon Rank Sum tests were used for comparisons of regular hubs and hyperinteracting hubs and for differential hub RNA-seq/ATAC-seq/H3K27ac ChIP-seq enrichment analyses with the wilcox.test function in R (version 4.2.1). Comparisons of RNA-seq differential gene enrichment between DND41/Rec-1 drug-sensitive/drug-resistant cells for selected genes were evaluated with Student’s t-tests (n = 3). Finally, statistical values for the comparison of probe distances in cumulative distribution plots were calculated using a Kolmogorov-Smirnov test. All p values less than 1E-5 were rounded up one decimal place (e.g. p = 8E-7 becomes p = 1E-6) for reporting in the text and figures. Unless otherwise noted, all box and whisker plots are shown without outliers to facilitate data visualization.

### Authors Contributions

Conceptualization: R.B.F., B.P.; Methodology: B.P., N.B., R.B.F., G.W.S., Y.Z.; Investigation: B.P., N.B., Y.Z., J.P., R.B.F.; Formal Analysis: B.P., N.B., Y.Z., J.P.; Resources and Reagents: R.B.F., Z.M.; Writing-Original Draft: B.P., R.B.F.; Writing-Review & Editing: Y.Z., N.B., J.P., G.W.S.; Z.M.; Funding Acquisition: R.B.F., Z.M., Supervision: R.B.F.

## Acknowledgments

We thank Golnaz Vahedi and her laboratory for their helpful discussion and support. This work was supported by R01-CA248041 and R01-CA230800 to R.B.F.

## Data Availability

All data needed to evaluate the conclusions in the paper are present in the paper and/or the Supplementary Materials. Newly generated sequencing datasets are available through GEO (GSE268228).

## Code Availability

All the code needed to evaluate the conclusions in the paper available at https://github.com/faryabiLab/hierarchical-spectral-clustering and https://github.com/faryabiLab/dockerize-workflows/tree/master/workflows.

