## Supplementary Figures for "Enhancer-promoter hubs organize transcriptional networks promoting oncogenesis and drug resistance"

**Affiliations:**
1 Department of Pathology and Laboratory Medicine, University of Pennsylvania, Philadelphia, PA, USA
2 Penn Epigenetics Institute, University of Pennsylvania, Philadelphia, PA, USA
3 Abramson Family Cancer Research Institute, University of Pennsylvania, Philadelphia, PA, USA
4 Genentech, South San Francisco, CA, USA
5 Princess Margaret Cancer Center, University Health Network, Toronto, ON, Canada

### Figure S1

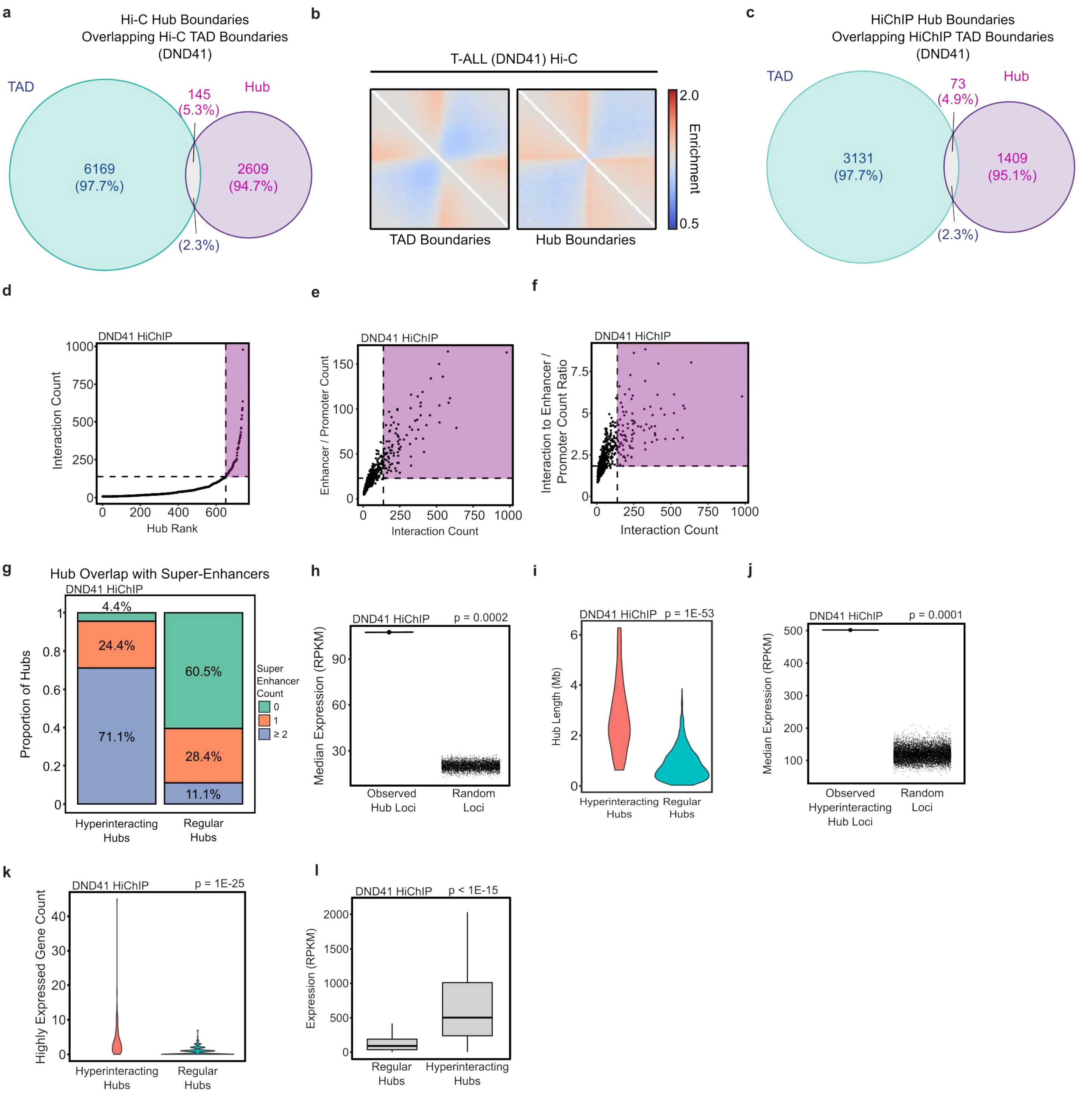

a
Hi-C Hub Boundaries
Overlapping Hi-C TAD Boundaries
(DND41)
TAD
Hub
145
(5.3%)
6169
(97.7%)
2609
(94.7%)
(2.3%)
b
T-ALL (DND41) Hi-C
2.0
Enrichment
0.5
TAD Boundaries
Hub Boundaries
c
HiChIP Hub Boundaries
Overlapping HiChIP TAD Boundaries
(DND41)
TAD
Hub
73
(4.9%)
3131
(97.7%)
1409
(95.1%)
(2.3%)
d
DND41 HiChIP
Interaction Count
Hub Rank
e
DND41 HiChIP
Enhancer / Promoter Count
Interaction Count
f
DND41 HiChIP
Interaction to Enhancer / Promoter Count Ratio
Interaction Count
g
Hub Overlap with Super-Enhancers
DND41 HiChIP
Proportion of Hubs
4.4%
24.4%
71.1%
60.5%
28.4%
11.1%
Super Enhancer Count
0
1
≥ 2
Hyperinteracting Hubs
Regular Hubs
h
DND41 HiChIP
p = 0.0002
Median Expression (RPKM)
Observed Hub Loci
Random Loci
i
DND41 HiChIP
p = 1E-53
Hub Length (Mb)
Hyperinteracting Hubs
Regular Hubs
j
DND41 HiChIP
p = 0.0001
Median Expression (RPKM)
Observed Hyperinteracting Hub Loci
Random Loci
k
DND41 HiChIP
p = 1E-25
Highly Expressed Gene Count
Hyperinteracting Hubs
Regular Hubs
l
DND41 HiChIP
p < 1E-15
Expression (RPKM)
Regular Hubs
Hyperinteracting Hubs

**Figure S1: SMC1 HiChIP corroborates Hi-C data and identifies T-ALL enhancer-promoter hubs.**

a: Venn diagram showing counts and percentages of overlap of enhancer-promoter hub and TAD boundaries detected from Hi-C of T-ALL DND41 cells.

b: Pile-up plots of the +/- 250 Kb surrounding Hi-C TAD (left) and Hi-C hub (right) boundaries demonstrating the relative strength of each type of boundary in terms of insulation score from DND41 Hi-C.

c: Venn diagram showing counts and percentages of overlap of enhancer-promoter hub and TAD boundaries detected from SMC1 HiChIP of T-ALL DND41 cells.

d: Enhancer-promoter hubs detected from T-ALL DND41 SMC1 HiChIP are plotted in ascending order of their total interactivity. Similar to Hi-C data, ranking of enhancer-promoter hubs' total interactivity detected by SMC1 HiChIP reveals asymmetrical distribution of interactions among enhancers and promoters across the T-ALL genome, and delineates two classes of regular and hyperinteracting hubs. Hyperinteracting hubs are marked within the purple region and defined as the hubs above the elbow of the total interactivity ranking.

e: Plotting SMC1 HiChIP interaction count vs. enhancer/promoter element count in each T-ALL DND41 hub demonstrates that the number of interactions increases concordantly with the number of enhancer-promoter elements in enhancer-promoter hubs. The purple region marks hyperinteracting hubs.

f: Plotting SMC1 HiChIP interaction count vs. ratio of SMC1 HiChIP interaction to enhancer/promoter counts shows that the T-ALL DND41 enhancer-promoter hubs with more interactions have more interconnected enhancer-promoter nodes, further supporting their hyperinteractivity. The purple region marks hyperinteracting hubs.

g: Stacked bar plots showing percentage of SMC1 HiChIP hyperconnected and regular hubs that overlap with 0, 1, or greater than or equal to 2 super-enhancers in T-ALL DND41.

h: T-ALL DND41 SMC1 HiChIP hubs are markedly enriched for transcriptional activity. Median expression of genes within loci of DND41 SMC1 HiChIP hubs is compared with a permutation model as indicated by black points. Each black point corresponds to median gene expression observed in an iteration of the permutation model, which is comprised of 5,000 sets of randomly selected genomic regions with chromosomes, genomic lengths, and loci counts matching those of the DND41 SMC1 HiChIP hubs. P-value: empirical p-value.

i: Violin plots showing distribution of genomic lengths of SMC1 HiChIP hyperinteracting and regular hubs in T-ALL DND41. P-value: two-tailed Wilcoxon rank sum test.

j: T-ALL DND41 SMC1 HiChIP hyperinteracting hubs are markedly enriched for transcriptional activity. Median expression of genes within loci of DND41 SMC1 HiChIP hyperinteracting hubs is compared with a permutation model as indicated by black points. Each black point corresponds to median gene expression observed in an iteration of the permutation model, which is comprised of 10,000 sets of randomly selected genomic regions with chromosomes, genomic lengths, and loci counts matching those of the DND41 SMC1 HiChIP hyperinteracting hubs. P-value: empirical p-value.

k: Violin plots showing distribution of the number of highly expressed genes within SMC1 HiChIP hyperinteracting and regular hubs in T-ALL DND41. Highly expressed genes are defined as those in the top 2.5% quantile of gene expression. P-value: two-tailed Wilcoxon rank sum test.

l: Box-and-whisker plots comparing transcription levels of T-ALL DND41 SMC1 HiChIP regular and hyperinteracting hubs show that hyperinteracting hubs are significantly more transcribed than regular hubs (without genomic span normalization). Box-and-whisker plots: center line, median; box limits, upper (75th) and lower (25th) percentiles; whiskers, 1.53 interquartile range. P-value: two-tailed Wilcoxon rank sum test.

### Figure S2

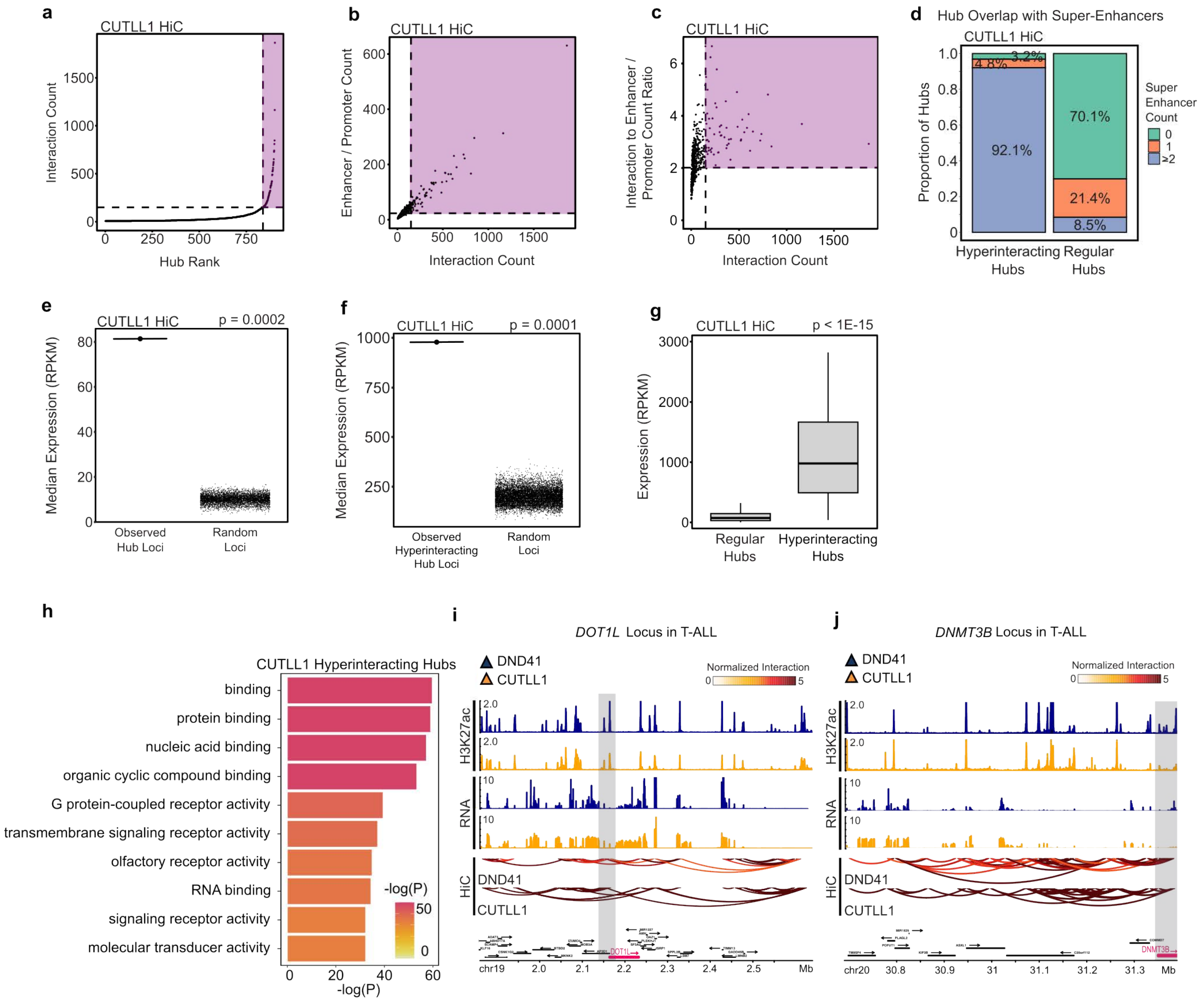
a
CUTLL1 HiC
Interaction Count
Hub Rank
b
CUTLL1 HiC
Enhancer / Promoter Count
Interaction Count
c
CUTLL1 HiC
Interaction to Enhancer / Promoter Count Ratio
Interaction Count
d
Hub Overlap with Super-Enhancers
CUTLL1 HiC
Proportion of Hubs
92.1%
4.8%
3.2%
70.1%
21.4%
8.5%
Super Enhancer Count
0
1
≥2
Hyperinteracting Hubs
Regular Hubs
e
CUTLL1 HiC
p = 0.0002
Median Expression (RPKM)
Observed Hub Loci
Random Loci
f
CUTLL1 HiC
p = 0.0001
Median Expression (RPKM)
Observed Hyperinteracting Hub Loci
Random Loci
g
CUTLL1 HiC
p < 1E-15
Expression (RPKM)
Regular Hubs
Hyperinteracting Hubs
h
CUTLL1 Hyperinteracting Hubs
binding
protein binding
nucleic acid binding
organic cyclic compound binding
G protein-coupled receptor activity
transmembrane signaling receptor activity
olfactory receptor activity
RNA binding
signaling receptor activity
molecular transducer activity
-log(P)
i
DOT1L Locus in T-ALL
DND41
CUTLL1
Normalized Interaction
H3K27ac
RNA
HiC
chr19
Mb
j
DNMT3B Locus in T-ALL
DND41
CUTLL1
Normalized Interaction
H3K27ac
RNA
HiC
chr20
Mb

**Figure S2: Similar to DND41, analysis of CUTLL1 shows that T-ALL hyperinteracting hubs are markedly transcribed and organize expression of genes encoding transcription factors.**

a: Enhancer-promoter hubs detected from T-ALL CUTLL1 Hi-C are plotted in ascending order of their total interactivity. Ranking of enhancer-promoter hubs' total interactivity detected by Hi-C reveals asymmetrical distribution of interactions among enhancers and promoters across the T-ALL genome, and delineates two classes of regular and hyperinteracting hubs. Hyperinteracting hubs are marked within the purple region and defined as the hubs above the elbow of the total interactivity ranking.

b: Plotting Hi-C interaction count vs. enhancer/promoter element count in each T-ALL CUTLL1 hub demonstrates that the number of interactions increases concordantly with the number of enhancer-promoter elements in enhancer-promoter hubs. The purple region marks hyperinteracting hubs.

c: Plotting Hi-C interaction count vs. ratio of Hi-C interaction to enhancer/promoter counts shows that the T-ALL CUTLL1 enhancer-promoter hubs with more interactions have more interconnected enhancer-promoter nodes, further supporting their hyperinteractivity. The purple region marks hyperinteracting hubs.

d: Stacked bar plots showing percentage of Hi-C hyperconnected and regular enhancer-promoter hubs that overlap with 0, 1, or greater than or equal to 2 super-enhancers in T-ALL CUTLL1.

e: T-ALL CUTLL1 Hi-C hubs are markedly enriched for transcriptional activity. Median expression of genes within loci of CUTLL1 Hi-C hubs is compared with a permutation model as indicated by black points. Each black point corresponds to median gene expression observed in an iteration of the permutation model, which is comprised of 5,000 sets of randomly selected genomic regions with chromosomes, genomic lengths, and loci counts matching those of the CUTLL1 Hi-C hubs. P-value: empirical p-value.

f: T-ALL CUTLL1 Hi-C hyperinteracting hubs are markedly enriched for transcriptional activity. Median expression of genes within loci of CUTLL1 Hi-C hyperinteracting hubs is compared with a permutation model as indicated by black points. Each black point corresponds to median gene expression observed in an iteration of the permutation model, which is comprised of 10,000 sets of randomly selected genomic regions with chromosomes, genomic lengths, and loci counts matching those of the CUTLL1 Hi-C hyperinteracting hubs. P-value: empirical p-value.

g: Box-and-whisker plots comparing transcription levels of T-ALL CUTLL1 Hi-C regular and hyperinteracting hubs show that hyperinteracting hubs are significantly more transcribed than regular hubs (without genomic span normalization). Box-and-whisker plots: center line, median; box limits, upper (75th) and lower (25th) percentiles; whiskers, 1.53 interquartile range. P-value: two-tailed Wilcoxon ranked sum test.

h: Gene ontology (GO) enrichment analysis showing that activities characteristic of transcription factors and cofactors are among the top 10 most enriched molecular functions associated with expressed genes located in T-ALL CUTLL1 Hi-C hyperinteracting hubs.

i, j: *DOT1L* and *DNMT3B* are located within hyperinteracting hubs in T-ALL. Hi-C arcs show that the gray box-marked *DOT1L* (i) and *DNMT3B* (j) form hyperinteracting enhancer-promoter hubs with several active regulatory elements and genes marked with H3K27ac ChIP-seq and RNA-seq, respectively, in T-ALL DND41 and CUTLL1. Bottom tracks indicate Ensembl gene position.

### Figure S3

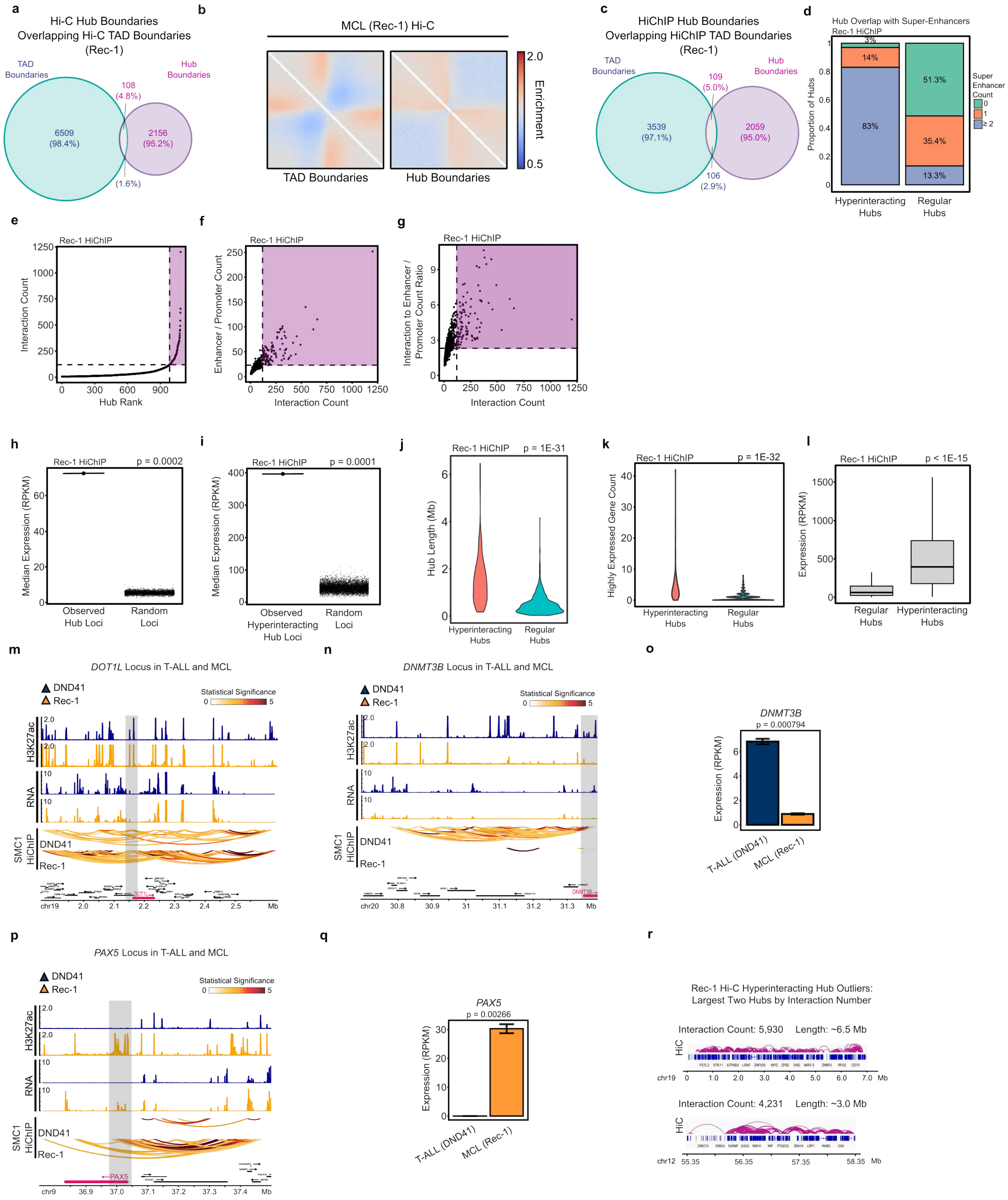

a
Hi-C Hub Boundaries Overlapping Hi-C TAD Boundaries (Rec-1)
TAD Boundaries
Hub Boundaries
6509 (98.4%)
108 (4.8%)
2156 (95.2%)
(1.6%)
b
MCL (Rec-1) Hi-C
TAD Boundaries
Hub Boundaries
Enrichment
2.0
0.5
c
HiChIP Hub Boundaries Overlapping HiChIP TAD Boundaries (Rec-1)
TAD Boundaries
Hub Boundaries
3539 (97.1%)
109 (5.0%)
2059 (95.0%)
106 (2.9%)
d
Hub Overlap with Super-Enhancers
Rec-1 HiChIP
Proportion of Hubs
Super Enhancer Count
0
1
≥ 2
3%
14%
83%
51.3%
35.4%
13.3%
Hyperinteracting Hubs
Regular Hubs
e
Rec-1 HiChIP
Interaction Count
Hub Rank
f
Rec-1 HiChIP
Enhancer / Promoter Count
Interaction Count
g
Rec-1 HiChIP
Interaction to Enhancer / Promoter Count Ratio
Interaction Count
h
Rec-1 HiChIP
p = 0.0002
Median Expression (RPKM)
Observed Hub Loci
Random Loci
i
Rec-1 HiChIP
p = 0.0001
Median Expression (RPKM)
Observed Hyperinteracting Hub Loci
Random Loci
j
Rec-1 HiChIP
p = 1E-31
Hub Length (Mb)
Hyperinteracting Hubs
Regular Hubs
k
Rec-1 HiChIP
p = 1E-32
Highly Expressed Gene Count
Hyperinteracting Hubs
Regular Hubs
l
Rec-1 HiChIP
p < 1E-15
Expression (RPKM)
Regular Hubs
Hyperinteracting Hubs
m
DOT1L Locus in T-ALL and MCL
DND41
Rec-1
Statistical Significance
H3K27ac
RNA
SMC1 HiChIP
DOT1L
chr19
Mb
n
DNMT3B Locus in T-ALL and MCL
DND41
Rec-1
Statistical Significance
H3K27ac
RNA
SMC1 HiChIP
DNMT3B
chr20
Mb
o
DNMT3B
p = 0.000794
Expression (RPKM)
T-ALL (DND41)
MCL (Rec-1)
p
PAX5 Locus in T-ALL and MCL
DND41
Rec-1
Statistical Significance
H3K27ac
RNA
SMC1 HiChIP
PAX5
chr9
Mb
q
PAX5
p = 0.00266
Expression (RPKM)
T-ALL (DND41)
MCL (Rec-1)
r
Rec-1 Hi-C Hyperinteracting Hub Outliers: Largest Two Hubs by Interaction Number
Interaction Count: 5,930 Length: ~6.5 Mb
HiC
chr19
Mb
Interaction Count: 4,231 Length: ~3.0 Mb
HiC
chr12
Mb

**Figure S3: SMC1 HiChIP corroborates Hi-C data and identifies MCL enhancer-promoter hubs.**

a: Venn diagram showing counts and percentages of overlap of enhancer-promoter hub and TAD boundaries detected from Hi-C of MCL Rec-1 cells.

b: Pile-up plots of the +/- 250 Kb surrounding Hi-C TAD (left) and Hi-C hub (right) boundaries demonstrating the relative strength of each type of boundary in terms of insulation score from Rec-1 Hi-C.

c: Venn diagram showing counts and percentages of overlap of enhancer-promoter hub and TAD boundaries detected from SMC1 HiChIP of MCL Rec-1 cells.

d: Stacked bar plots showing percentage of SMC1 HiChIP hyperconnected and regular hubs that overlap with 0, 1, or greater than or equal to 2 super-enhancers in MCL Rec-1.

e: Enhancer-promoter hubs detected from MCL Rec-1 SMC1 HiChIP are plotted in ascending order of their total interactivity. Similar to Hi-C data, ranking of enhancer-promoter hubs' total interactivity detected by SMC1 HiChIP reveals asymmetrical distribution of interactions among enhancers and promoters across the MCL genome, and delineates two classes of regular and hyperinteracting hubs. Hyperinteracting hubs are marked within the purple region and defined as the hubs above the elbow of the total interactivity ranking.

f: Plotting SMC1 HiChIP interaction count vs. enhancer/promoter element count in each MCL Rec-1 hub demonstrates that the number of interactions increases concordantly with the number of enhancer-promoter elements in enhancer-promoter hubs. The purple region marks hyperinteracting hubs.

g: Plotting SMC1 HiChIP interaction count vs. ratio of SMC1 HiChIP interaction to enhancer/promoter counts shows that the MCL Rec-1 enhancer-promoter hubs with more interactions have more interconnected enhancer-promoter nodes, further supporting their hyperinteractivity. The purple region marks hyperinteracting hubs.

h: MCL Rec-1 SMC1 HiChIP hubs are markedly enriched for transcriptional activity. Median expression of genes within loci of Rec-1 SMC1 HiChIP hubs is compared with a permutation model as indicated by black points. Each black point corresponds to median gene expression observed in an iteration of the permutation model, which is comprised of 5,000 sets of randomly selected genomic regions with chromosomes, genomic lengths, and loci counts matching those of the Rec-1 SMC1 HiChIP hubs. P-value: empirical p-value.

i: MCL Rec-1 SMC1 HiChIP hyperinteracting hubs are markedly enriched for transcriptional activity. Median expression of genes within loci of Rec-1 SMC1 HiChIP hyperinteracting hubs is compared with a permutation model as indicated by black points. Each black point corresponds to median gene expression observed in an iteration of the permutation model, which is comprised of 10,000 sets of randomly selected genomic regions with chromosomes, genomic lengths, and loci counts matching those of the Rec-1 SMC1 HiChIP hyperinteracting hubs. P-value: empirical p-value.

j: Violin plots showing distribution of genomic lengths of SMC1 HiChIP hyperinteracting and regular hubs in MCL Rec-1. P-value: two-tailed Wilcoxon rank sum test.

k: Violin plots showing distribution of the number of highly expressed genes within SMC1 HiChIP hyperinteracting and regular hubs in MCL Rec-1. Highly expressed genes are defined as those in the top 2.5% quantile of gene expression. P-value: two-tailed Wilcoxon rank sum test.

l: Box-and-whisker plots comparing transcription levels of MCL Rec-1 SMC1 HiChIP regular and hyperinteracting hubs show that hyperinteracting hubs are significantly more transcribed than regular hubs (without genomic span normalization) Box-and-whisker plots: center line, median; box limits, upper (75th) and lower (25th) percentiles; whiskers, 1.53 interquartile range. P-value: two-tailed Wilcoxon ranked sum test.

m: *DOT1L* is located in a hyperinteracting hub in T-ALL and MCL. SMC1 HiChIP arcs show that the gray box-marked *DOT1L* forms a hyperinteracting enhancer-promoter hub with several active regulatory elements and genes marked with H3K27ac ChIP-seq and RNA-seq, respectively, in T-ALL DND41 and MCL Rec-1. Bottom tracks indicate Ensembl gene position.

n: *DNMT3B* is located in a hyperinteracting hub in T-ALL but not MCL. SMC1 HiChIP arcs show that the gray box-marked *DNMT3B* forms a hyperinteracting enhancer-promoter hub with several active regulatory elements and genes marked with H3K27ac ChIP-seq and RNA-seq, respectively, in T-ALL DND41 but not MCL Rec-1. Bottom tracks indicate Ensembl gene position.

o: *DNMT3B* is significantly more expressed in T-ALL DND41 than MCL Rec-1. Barplots of normalized RNA-seq reads showing *DNMT3B* mRNA levels in DND41 and Rec-1 cells. N: 3; P-value: unpaired student t-test; error bars: +/- 2 SE.

p: *PAX5* is located in a hyperinteracting hub in MCL but not T-ALL. SMC1 HiChIP arcs show that the gray box-marked *PAX5* forms a hyperinteracting enhancer-promoter hub with several active regulatory elements and genes marked with H3K27ac ChIP-seq and RNA-seq, respectively, in MCL Rec-1 but not T-ALL DND41. Bottom tracks indicate Ensembl gene position.

q: *PAX5* is significantly more expressed in MCL Rec-1 than T-ALL DND41. Barplots of normalized RNA-seq reads showing *PAX5* mRNA levels in DND41 and Rec-1 cells. N: 3; P-value: unpaired student t-test; error bars: +/- 2 SE.

r: Genomic tracks of MCL Rec-1 Hi-C contacts over the two outlier hyperinteracting hubs show that these loci contain several adjacent, overlapping clusters. These large outlier hubs skew calculation of the hyperinteracting hub interaction cutoff and contribute to the lower level of similarity between Rec-1 Hi-C and SMC1 HiChIP hyperinteracting hubs seen in Fig. 3i.

### Figure S4

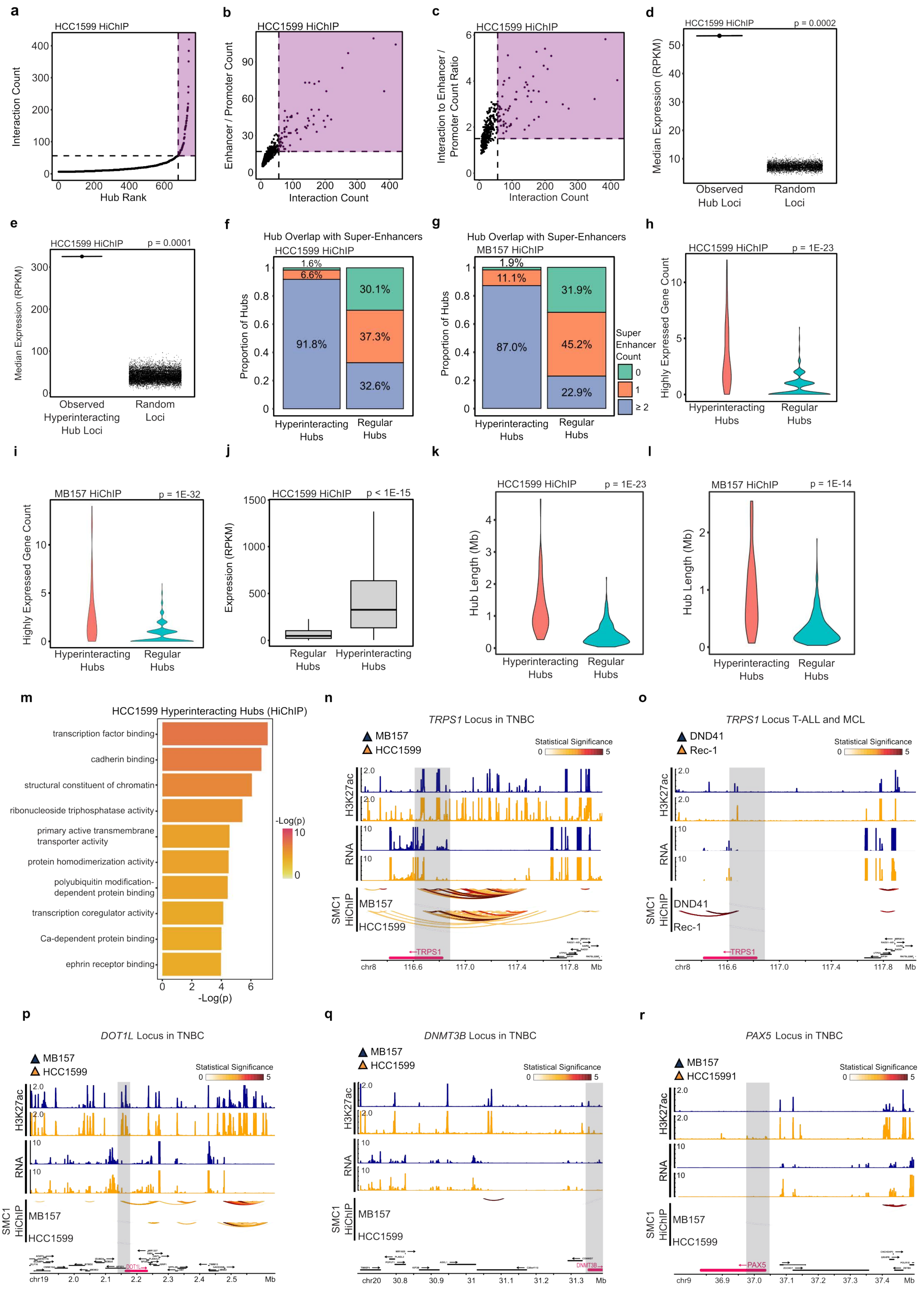
a
HCC1599 HiChIP
Interaction Count
Hub Rank
b
HCC1599 HiChIP
Enhancer / Promoter Count
Interaction Count
c
HCC1599 HiChIP
Interaction to Enhancer / Promoter Count Ratio
Interaction Count
d
HCC1599 HiChIP
p = 0.0002
Median Expression (RPKM)
Observed Hub Loci
Random Loci
e
HCC1599 HiChIP
p = 0.0001
Median Expression (RPKM)
Observed Hyperinteracting Hub Loci
Random Loci
f
Hub Overlap with Super-Enhancers
HCC1599 HiChIP
Proportion of Hubs
1.6%
6.6%
91.8%
30.1%
37.3%
32.6%
Hyperinteracting Hubs
Regular Hubs
g
Hub Overlap with Super-Enhancers
MB157 HiChIP
Proportion of Hubs
1.9%
11.1%
87.0%
31.9%
45.2%
22.9%
Hyperinteracting Hubs
Regular Hubs
Super Enhancer Count
0
1
≥ 2
h
HCC1599 HiChIP
p = 1E-23
Highly Expressed Gene Count
Hyperinteracting Hubs
Regular Hubs
i
MB157 HiChIP
p = 1E-32
Highly Expressed Gene Count
Hyperinteracting Hubs
Regular Hubs
j
HCC1599 HiChIP
p < 1E-15
Expression (RPKM)
Regular Hubs
Hyperinteracting Hubs
k
HCC1599 HiChIP
p = 1E-23
Hub Length (Mb)
Hyperinteracting Hubs
Regular Hubs
l
MB157 HiChIP
p = 1E-14
Hub Length (Mb)
Hyperinteracting Hubs
Regular Hubs
m
HCC1599 Hyperinteracting Hubs (HiChIP)
transcription factor binding
cadherin binding
structural constituent of chromatin
ribonucleoside triphosphatase activity
primary active transmembrane transporter activity
protein homodimerization activity
polyubiquitin modification-dependent protein binding
transcription coregulator activity
Ca-dependent protein binding
ephrin receptor binding
-Log(p)
n
TRPS1 Locus in TNBC
MB157
HCC1599
Statistical Significance
H3K27ac
RNA
SMC1 HiChIP
TRPS1
chr8
Mb
o
TRPS1 Locus T-ALL and MCL
DND41
Rec-1
Statistical Significance
H3K27ac
RNA
SMC1 HiChIP
TRPS1
chr8
Mb
p
DOT1L Locus in TNBC
MB157
HCC1599
Statistical Significance
H3K27ac
RNA
SMC1 HiChIP
DOT1L
chr19
Mb
q
DNMT3B Locus in TNBC
MB157
HCC1599
Statistical Significance
H3K27ac
RNA
SMC1 HiChIP
DNMT3B
chr20
Mb
r
PAX5 Locus in TNBC
MB157
HCC15991
Statistical Significance
H3K27ac
RNA
SMC1 HiChIP
PAX5
chr9
Mb

**Figure S4: Similar to MB157, analysis of HCC1599 shows that TNBC hyperinteracting hubs are markedly transcribed and organize expression of genes encoding transcription factors.**

a: Enhancer-promoter hubs detected from TNBC HCC1599 SMC1 HiChIP are plotted in ascending order of their total interactivity. Ranking of enhancer-promoter hubs' total interactivity confirms MB157 data and shows asymmetrical distribution of interactions among enhancers and promoters across the TNBC genome, delineating two classes of regular and hyperinteracting hubs. Hyperinteracting hubs are marked within the purple region and defined as the hubs above the elbow of the total interactivity ranking.

b: Plotting SMC1 HiChIP interaction count vs. enhancer/promoter element count in each TNBC HCC1599 hub demonstrates that the number of interactions increases concordantly with the number of enhancer-promoter elements in enhancer-promoter hubs. The purple region marks hyperinteracting hubs.

c: Plotting SMC1 HiChIP interaction count vs. ratio of SMC1 HiChIP interaction to enhancer/promoter counts shows that similar to MB157, TNBC HCC1599 enhancer-promoter hubs with more interactions have more interconnected enhancer-promoter nodes, further supporting their hyperinteractivity. The purple region marks hyperinteracting hubs.

d: TNBC HCC1599 hubs are markedly enriched for transcriptional activity. Median expression of genes within loci of HCC1599 hubs is compared with a permutation model as indicated by black points. Each black point corresponds to median gene expression observed in an iteration of the permutation model, which is comprised of 5,000 sets of randomly selected genomic regions with chromosomes, genomic lengths, and loci counts matching those of the HCC1599 hubs. P-value: empirical p-value.

e: TNBC HCC1599 hyperinteracting hubs are markedly enriched for transcriptional activity. Median expression of genes within loci of HCC1599 hyperinteracting hubs is compared with a permutation model as indicated by black points. Each black point corresponds to median gene expression observed in an iteration of the permutation model, which is comprised of 10,000 sets of randomly selected genomic regions with chromosomes, genomic lengths, and loci counts matching those of the HCC1599 hyperinteracting hubs. P-value: empirical p-value.

f: Stacked bar plots showing percentage of HCC1599 SMC1 HiChIP hyperconnected and regular hubs that overlap with 0, 1, or greater than or equal to 2 super-enhancers in TNBC HCC1599.

g: Stacked bar plots showing percentage of MB157 SMC1 HiChIP hyperconnected and regular hubs that overlap with 0, 1, or greater than or equal to 2 super-enhancers in TNBC MB157.

h: Violin plots showing distribution of the number of highly expressed genes within SMC1 HiChIP hyperinteracting and regular hubs in TNBC HCC1599. Highly expressed genes are defined as those in the top 2.5% quantile of gene expression. P-value: two-tailed Wilcoxon rank sum test.

i: Violin plots showing distribution of the number of highly expressed genes within SMC1 HiChIP hyperinteracting and regular hubs in TNBC MB157. Highly expressed genes are defined as those in the top 2.5% quantile of gene expression. P-value: two-tailed Wilcoxon rank sum test.

j: Box-and-whisker plots comparing transcription levels of TNBC HCC1599 SMC1 HiChIP regular and hyperinteracting hubs show that hyperinteracting hubs are significantly more transcribed than regular hubs (without genomic distance normalization). Box-and-whisker plots: center line, median; box limits, upper (75th) and lower (25th) percentiles; whiskers, 1.53 interquartile range. P-value: two-tailed Wilcoxon rank sum test.

k: Violin plots showing distribution of genomic lengths of SMC1 HiChIP hyperinteracting and regular hubs in TNBC HCC1599. P-value: two-tailed Wilcoxon rank sum test.

l: Violin plots showing distribution of genomic lengths of SMC1 HiChIP hyperinteracting and regular hubs in TNBC MB157. P-value: two-tailed Wilcoxon rank sum test.

m: Gene ontology (GO) enrichment analysis showing that similar to MB157, activities characteristic of transcription factors and cofactors are among the top 10 most enriched molecular functions associated with expressed genes located in TNBC HCC1599 hyperinteracting hubs.

n, o: *TRPS1* is located in a hyperinteracting hub in TNBC but not T-ALL and MCL. SMC1 HiChIP arcs show that the gray box-marked *TRPS1* forms a hyperinteracting enhancer-promoter hub with several active regulatory elements and genes marked with H3K27ac ChIP-seq and RNA-seq, respectively, in TNBC MB157 and HCC1599 (n) but not T-ALL DND41 and MCL Rec-1 (o). Bottom tracks indicate Ensembl gene position.

p, q, r: *DOT1L*, *DNMT3B*, and *PAX5* are not located in hyperinteracting hubs in TNBC. SMC1 HiChIP arcs show that the gray box-marked *DOT1L* (p), *DNMT3B* (q), and *PAX5* (r) do not form hyperinteracting enhancer-promoter hubs in TNBC MB157 and HCC1599. Bottom tracks indicate Ensembl gene position.

### Figure S5

a

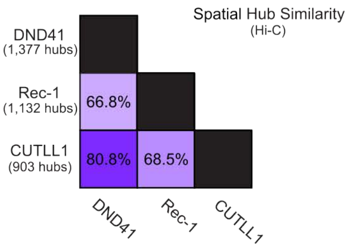


b

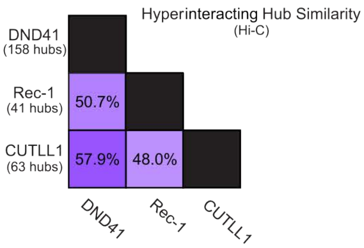


c

| Chromosome | Start (MB) | End (MB) | Selected Expressed Genes* |
|---|---|---|---|
| chr1 | 153.51 | 156.83 | EFNA4, PYGO2, SHC1, etc. |
| chr1 | 234.30 | 236.03 | ARID4B, TOMM20, IRF2BP2, etc. |
| chr6 | 25.99 | 29.72 | HIST1H1B, HIST1H4K, HIST1H2BK, etc. |
| chr6 | 29.89 | 36.16 | HLA-E, ATAT1, ABCF1, DDR1, etc. |
| chr7 | 98.27 | 100.97 | TRIP6, PLOD3, etc. |
| chr8 | 128.75 | 130.69 | MYC, etc. |
| chr8 | 144.25 | 146.29 | PARP10, MAF1, ZNF16, PUF60, etc. |
| chr19 | 36.87 | 41.11 | SIRT2, NFKBIB, ACTN4, PSMD8, etc. |
| chr20 | 52.19 | 52.83 | ZNF217, etc. |
| chr21 | 42.69 | 47.40 | SIK1, HSF2BP, RRP1B, etc. |

*Hub genomic intervals and expressed genes are provided in terms of DND41 SMC1 HiChIP hubs

d

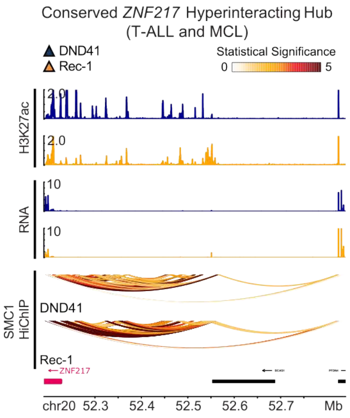


e

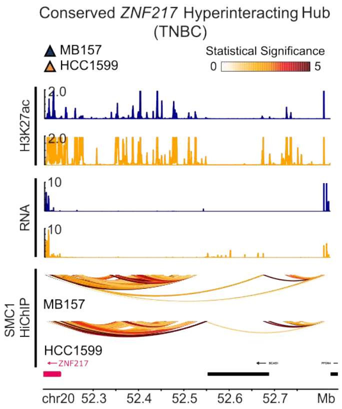

**Figure S5: Hyperinteracting enhancer-promoter hubs shared across T-ALL, MCL, and TNBC are located at genes involved in oncogenesis.**

a: Matrix of pairwise Hi-C hub similarity shows that these topological assemblies are somewhat conserved across T-ALL and MCL cells. Similarity between each pair is calculated as the percentage of enhancer-promoter hubs with overlapping genomic intervals. Total hub counts for each cell line is listed on the left.

b: Matrix of pairwise Hi-C hyperinteracting hub similarity shows that these topological assemblies are more lineage associated than all enhancer-promoter hubs in T-ALL and MCL cells. Similarity between each pair is calculated as the percentage of hyperinteracting hubs with overlapping genomic intervals. Total hyperinteracting hub counts for each cell line is listed on the left.

c: Table demonstrating the SMC1 HiChIP hyperinteracting hubs that are conserved across T-ALL, MCL, and TNBC on the basis of genomic overlap. Each hyperinteracting hub is annotated with selected expressed genes in DND41, several of which are related to cancer pathobiology. The genomic interval of each shared DND41 SMC1 HiChIP hyperinteracting hub is reported in the table.

d, e: The *ZNF217* hyperinteracting hub is present in T-ALL, MCL, and TNBC. SMC1 HiChIP arcs show that the gray box-marked *ZNF217* forms a hyperinteracting enhancer-promoter hub with several active regulatory elements and genes marked with H3K27ac ChIP-seq and RNA-seq, respectively, in T-ALL DND41 and MCL Rec-1 (d) as well as TNBC MB157 and HCC1599 (e). Bottom tracks indicate Ensembl gene position.

### Figure S6

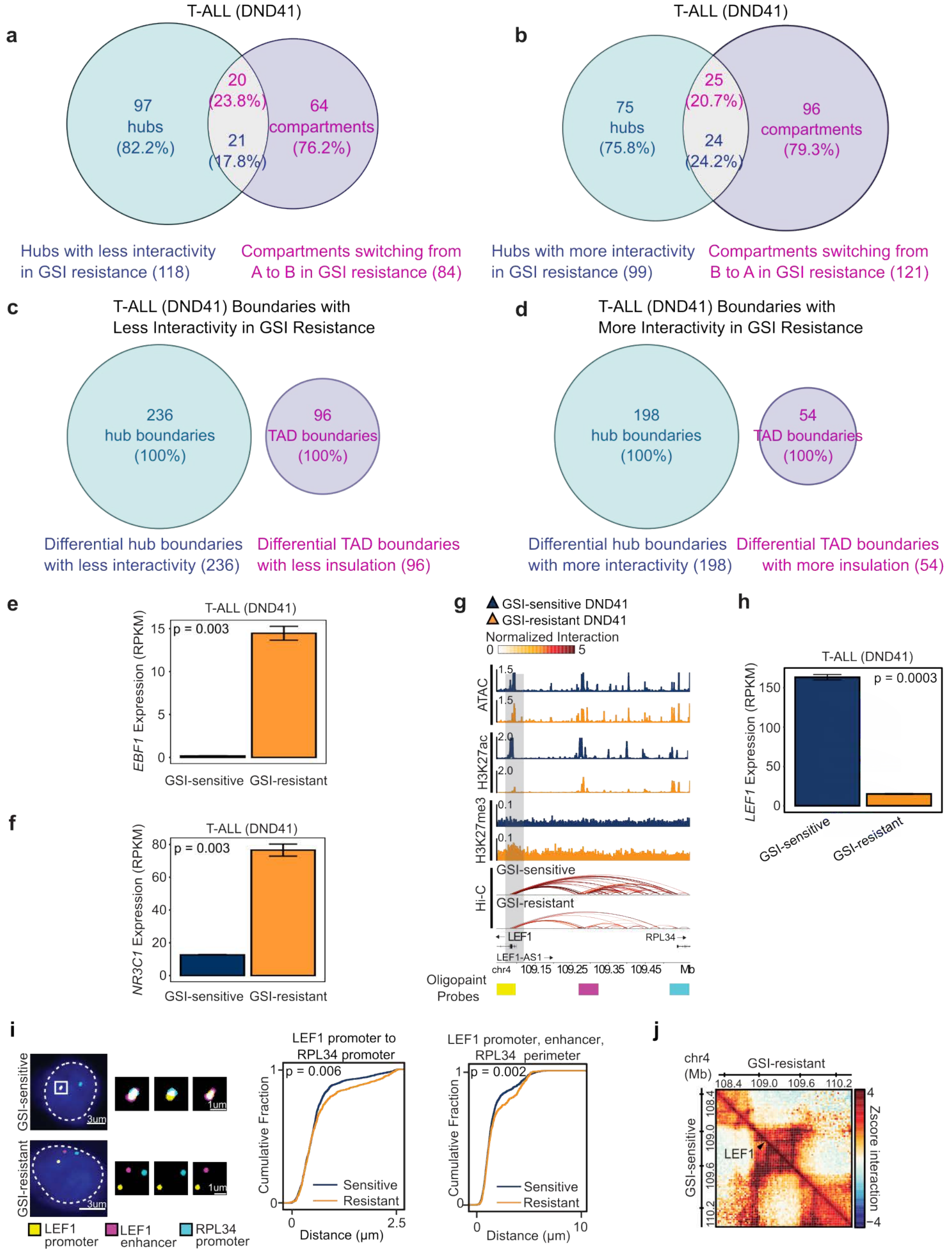

a
T-ALL (DND41)
97 hubs (82.2%)
20 (23.8%)
21 (17.8%)
64 compartments (76.2%)
Hubs with less interactivity in GSI resistance (118)
Compartments switching from A to B in GSI resistance (84)
b
T-ALL (DND41)
75 hubs (75.8%)
25 (20.7%)
24 (24.2%)
96 compartments (79.3%)
Hubs with more interactivity in GSI resistance (99)
Compartments switching from B to A in GSI resistance (121)
c
T-ALL (DND41) Boundaries with Less Interactivity in GSI Resistance
236 hub boundaries (100%)
96 TAD boundaries (100%)
Differential hub boundaries with less interactivity (236)
Differential TAD boundaries with less insulation (96)
d
T-ALL (DND41) Boundaries with More Interactivity in GSI Resistance
198 hub boundaries (100%)
54 TAD boundaries (100%)
Differential hub boundaries with more interactivity (198)
Differential TAD boundaries with more insulation (54)
e
T-ALL (DND41)
p = 0.003
EBF1 Expression (RPKM)
GSI-sensitive GSI-resistant
f
T-ALL (DND41)
p = 0.003
NR3C1 Expression (RPKM)
GSI-sensitive GSI-resistant
g
GSI-sensitive DND41
GSI-resistant DND41
Normalized Interaction
ATAC
H3K27ac
H3K27me3
Hi-C
GSI-sensitive
GSI-resistant
LEF1
RPL34
LEF1-AS1
chr4 109.15 109.25 109.35 109.45 Mb
Oligopaint Probes
h
T-ALL (DND41)
p = 0.0003
LEF1 Expression (RPKM)
GSI-sensitive GSI-resistant
i
GSI-sensitive
GSI-resistant
LEF1 promoter
LEF1 enhancer
RPL34 promoter
LEF1 promoter to RPL34 promoter
p = 0.006
Cumulative Fraction
Sensitive
Resistant
Distance (μm)
LEF1 promoter, enhancer, RPL34 perimeter
p = 0.002
Cumulative Fraction
Sensitive
Resistant
Distance (μm)
j
chr4 (Mb)
GSI-resistant
GSI-sensitive
LEF1
Zscore interaction

**Figure S6: Loss of *LEF1* hyperinteracting hub coincides with decrease in chromatin activity, gene expression, and architectural stripe in GSI-resistant T-ALL.**

a: Venn diagram showing counts and percentages of overlap of differential Hi-C hubs that markedly lost interactivity and compartments that switched from A to B in GSI-resistant T-ALL DND41.

b: Venn diagram showing counts and percentages of overlap of differential Hi-C hubs that markedly gained interactivity and compartments that switched from B to A in GSI-resistant T-ALL DND41.

c: Venn diagram showing counts and percentages of overlap of differential Hi-C hubs and TAD boundaries that markedly lost interactivity and insulation (respectively) in GSI-resistant T-ALL DND41.

d: Venn diagram showing counts and percentages of overlap of differential Hi-C hubs and TAD boundaries that markedly gained interactivity and insulation (respectively) in GSI-resistant T-ALL DND41.

e: *EBF1* is significantly upregulated in GSI-resistant T-ALL cells. Barplots of normalized RNA-seq reads showing *EBF1* mRNA levels in GSI-resistant and GSI-sensitive DND41. N: 3; P-value: unpaired student t-test; error bars: +/- 2 SE.

f: *NR3C1* is significantly upregulated in GSI-resistant T-ALL cells. Barplots of normalized RNA-seq reads showing *NR3C1* mRNA levels in GSI-resistant and GSI-sensitive DND41. N: 3; P-value: unpaired student t-test; error bars: +/- 2 SE.

g: Genome tracks at the *LEF1* locus show that loss of Hi-C interactions in the *LEF1* hyperinteracting hub in GSI-resistant DND41 cells coincide with marked decreases in chromatin accessibility and H3K27ac active chromatin mark. Oligopaint DNA FISH probes are marked with pseudo-color yellow (*LEF1* promoter), cyan (*RPL34* promoter), and magenta (*LEF1* enhancer) below the Ensembl gene track.

h: *LEF1* is significantly downregulated in GSI-resistant T-ALL cells. Barplots of normalized RNA-seq reads showing *LEF1* mRNA levels in GSI-resistant and GSI-sensitive DND41. N: 3; P-value: unpaired student t-test; error bars: +/- 2 SE.

i: *LEF1* hub expands in GSI-resistant T-ALL DND41 cells. Left: confocal microscopy images of representative cells and magnified images for comparison of 3-color *LEF1* promoter (yellow), *LEF1* enhancer (magenta), *RPL34* promoter (cyan) probes in GSI-sensitive (top) and GSI-resistant (bottom) cells are shown. Locations of the three 50 Kb Oligopaint probes for the *LEF1* promoter (yellow), *LEF1* enhancer (magenta), and *RPL34* promoter (cyan) are marked in (A). Center: cumulative distribution plots of the closest distance between the *LEF1* promoter and the *RPL34* promoter in the same cell are compared between 1,319 GSI-sensitive and 640 GSI-resistant allelic interactions. Mean (+/- S.D.) distance in GSI-sensitive and GSI-resistant cells is 0.643 (+/- 0.46) μm, and 0.742 (+/- 0.58) μm, respectively (Kolmogorov-Smirnov p-value = 0.006). Right: cumulative distribution plots of spatial perimeter formed by the *LEF1* promoter, *LEF1* enhancer, and *RPL34* promoter. Mean (+/- S.D.) in GSI-sensitive and GSI-resistant cells is 1.87 (+/- 1.26) μm, and 2.09 (+/- 1.40) μm, respectively (Kolmogorov-Smirnov p-value = 0.002).

j: Normalized Hi-C contact maps in GSI-resistant (upper triangle) and GSI-sensitive T-ALL DND41 cells (lower triangle) at the *LEF1* locus show that both loops and an architectural stripe interacting with arrowed-marked *LEF1* in GSI-sensitive DND41 are lost upon acquisition of GSI resistance.

### Figure S7

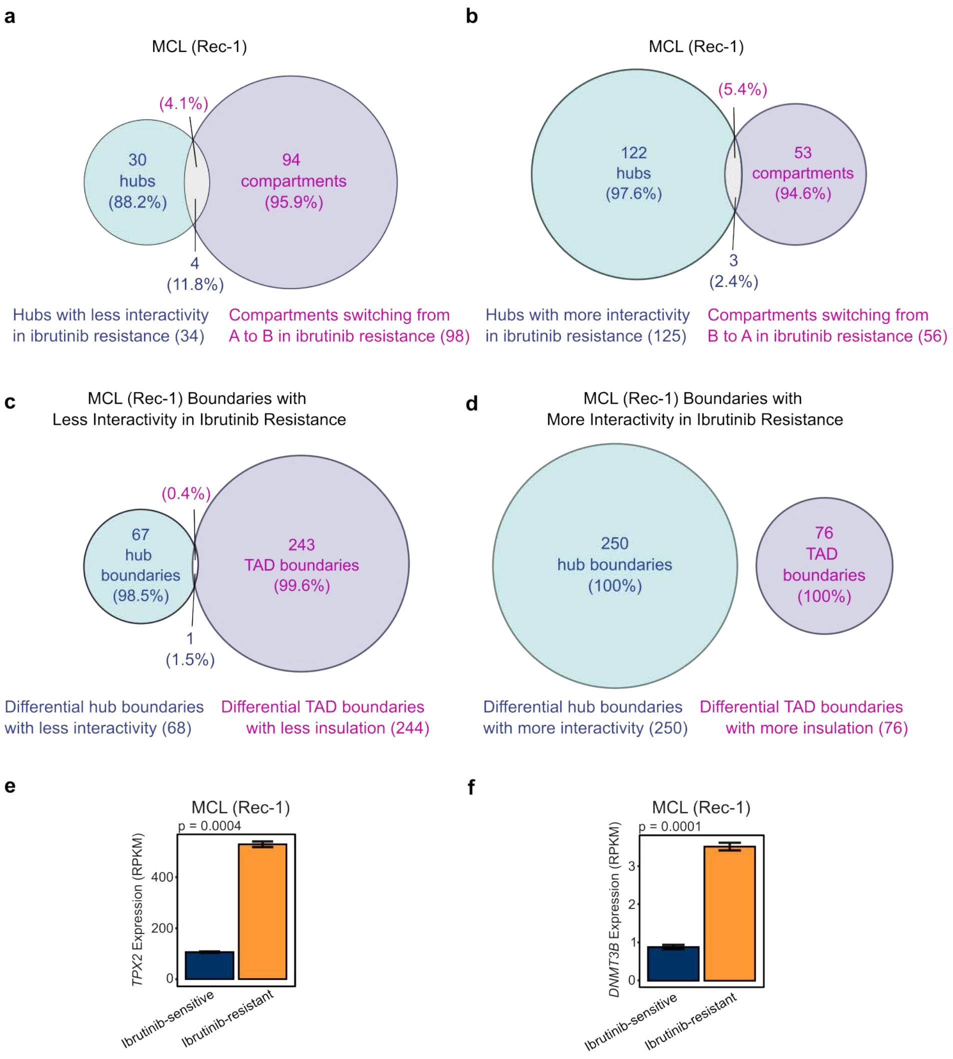

a
MCL (Rec-1)
30 hubs (88.2%)
(4.1%)
94 compartments (95.9%)
4 (11.8%)
Hubs with less interactivity in ibrutinib resistance (34)
Compartments switching from A to B in ibrutinib resistance (98)
b
MCL (Rec-1)
122 hubs (97.6%)
(5.4%)
53 compartments (94.6%)
3 (2.4%)
Hubs with more interactivity in ibrutinib resistance (125)
Compartments switching from B to A in ibrutinib resistance (56)
c
MCL (Rec-1) Boundaries with Less Interactivity in Ibrutinib Resistance
67 hub boundaries (98.5%)
(0.4%)
243 TAD boundaries (99.6%)
1 (1.5%)
Differential hub boundaries with less interactivity (68)
Differential TAD boundaries with less insulation (244)
d
MCL (Rec-1) Boundaries with More Interactivity in Ibrutinib Resistance
250 hub boundaries (100%)
76 TAD boundaries (100%)
Differential hub boundaries with more interactivity (250)
Differential TAD boundaries with more insulation (76)
e
MCL (Rec-1)
p = 0.0004
TPX2 Expression (RPKM)
0
200
400
Ibrutinib-sensitive
Ibrutinib-resistant
f
MCL (Rec-1)
p = 0.0001
DNMT3B Expression (RPKM)
0
1
2
3
Ibrutinib-sensitive
Ibrutinib-resistant

**Figure S7: Differential hubs are distinct from differential TADs and compartments between ibrutinib-sensitive and -resistant MCL Rec-1.**

a: Venn diagram showing counts and percentages of overlap of differential Hi-C hubs that markedly lost interactivity and compartments that switched from A to B in ibrutinib-resistant MCL Rec-1.

b: Venn diagram showing counts and percentages of overlap of differential Hi-C hubs that markedly gained interactivity and compartments that switched from B to A in ibrutinib-resistant MCL Rec-1.

c: Venn diagram showing counts and percentages of overlap of differential Hi-C hubs and TAD boundaries that markedly lost interactivity and insulation (respectively) in ibrutinib-resistant MCL Rec-1.

d: Venn diagram showing counts and percentages of overlap of differential Hi-C hubs and TAD boundaries that markedly gained interactivity and insulation (respectively) in ibrutinib-resistant MCL Rec-1.

e: *TPX2* is significantly upregulated in ibrutinib-resistant MCL Rec-1 cells. Barplots of normalized RNA-seq reads showing *TPX2* mRNA levels in ibrutinib-resistant and -sensitive Rec-1. N: 3; P-value: unpaired student t-test; error bars: +/- 2 SE. *TPX2* is a part of the strongly gained hyperinteracting hub containing *BCL2L1* that is illustrated in Fig. 7G.

f: *DNMT3B* is significantly upregulated in ibrutinib-resistant MCL Rec-1 cells. Barplots of normalized RNA-seq reads showing *DNMT3B* mRNA levels in ibrutinib-resistant and -sensitive Rec-1. N: 3; P-value: unpaired student t-test; error bars: +/- 2 SE. *DNMT3B* is a part of the strongly gained hyperinteracting hub containing *BCL2L1* that is illustrated in Fig. 7G.

**Supplemental Table Legends:**

**Table S1:** Annotated Hi-C and SMC1 HiChIP enhancer-promoter hubs for all cell lines and conditions.

**Table S2:** GO enrichment analysis output for expressed genes in all Hi-C and SMC1 HiChIP hyperinteracting enhancer-promoter hubs as well as for expressed genes in T-ALL DND41 and MCL Rec-1 differential hubs.

**Table S3**: Union and differential hubs identified between drug-sensitive and drug-resistant DND41 and Rec-1.

**Table S4**: EagleC structural rearrangement analysis output for Rec-1 ibrutinib-sensitive and ibrutinib-resistant Hi-C data.
